# From tumor microenvironment to immuno-therapeutic outcomes for solid tumors: A systems theoretic approach

**DOI:** 10.1101/2024.09.26.615155

**Authors:** Priyan Bhattacharya, Rajanikanth Vadigepalli

## Abstract

Increasing evidence suggests the tumor microenvironment (TME) governs solid tumor response to immune checkpoint inhibition (ICI). Decoding the relationship between the cell compositional diversity of the tumor microenvironment (TME) and the therapeutic outcomes has been a longstanding problem in solid tumor research. In this work, we develop a systems-theoretic formalism to decipher the key mechanisms of growth, proliferation, immune evasion, and drug resistance that run common across solid tumors in the context of Immune Checkpoint Inhibitors (ICI). We reconstructed a core TME network in common across most solid tumors, containing multiple tumor and non-tumor cell types in distinct functional states, and molecular agents mediating cellular signaling and cell-cell interactions. Our analysis shows that the core TME network is sufficient to yield a multiplicity of attractors corresponding to clinically observed TME subtypes namely, immune or fibro dominated, immune or fibro desert, and immune and fibro deficient. Importantly, the reachability around the pre-ICI attractors governs the response to ICI explaining the TME subtype-specific therapy outcomes. We analyzed the attractor transition network to identify subtype-specific combination therapies that can drive unresponsive TME to a responsive subtype. We derived mathematical conditions relating TME balances to determine the limits of the efficacy of combination therapies. Our results hold for a large class of smooth biochemical kinetics with monotone and bounded interactions and (semi-)concave proliferation rules. The analytical findings have been verified with extensive simulation of different TME sub-types. Overall, we propose a generalized systems formalism that accounts for the TME properties governing ICI response and can aid in designing intervention strategies for improved tumor prognosis.

**Significance Statement:** Non-responsive therapy outcomes have been a persistent problem in cancer treatment. Predicting the possibility of non-responsiveness to a particular therapy from the pre-treatment composition of the tumor microenvironment (TME) aids in designing appropriate combination treatment strategies toward an improved prognosis. The present work develops a systems-theoretic formalism that aims to unfold the mechanisms behind solid tumors’ growth, non-responsiveness, and recurrence. Unlike a single model, the proposed formalism does not assume any particular kinetics, barring some minimal assumptions. This enables us to explain some of the relevant observations made by the recent experimental studies. Finally, the closed-form conditions obtained for responsivity and recurrence can also guide novel therapeutic strategies that may be able to restore responsiveness to immunotherapy.

## 1 Introduction

Cell division and proliferation is a complex process involving several agents as flux modulators that aid in steering the cell population to a new metabolically compatible state [1]. Therefore, an investigation into the growth pattern of a particular cell type demands a holistic understanding of the surroundings that encompasses all possible *pro-* and *anti-*growth agents. It is conceivable that in the progression of cancer, which is characterized by *uncontrolled growth* at its core, the survival and growth of tumor cells are sustained by a number of surrounding non-tumor cells [2]. In this precise sense, the collection of tumor and non-tumor cells, along with the inter and intra-cellular interactions, constitute the underlying tumor microenvironment (TME) [3, 4]. The pro-tumor components aid in the growth, immune evasion, and lymph node metastasis, whereas the anti-tumoral components, mediated by the immune systems, promote killing and modulate the apoptosis rate of tumor cells [3]. Additionally, these pro- and anti-inflammatory modules also secrete cyto- and chemokines that can further trigger pro(anti)-tumor mechanisms [5]. It is to be noted that although the pro and anti-tumor attributes are mostly well-defined and distinct, it is not always possible to classify a particular module (cell type) in the TME vis-a-vis the pro-anti dichotomy [5]. For instance, despite being the main immune response driver, certain T cells can encourage tumor growth [6]. Therefore, it is necessary to determine the appropriate representation of the TME given the purpose of the study.

Depending on the proportion of different components in the TME, there can be several possible TME sub-types such as immune or fibro desert and fibro or immune-rich [3, 4, 7, 8]. These TME subtypes produce distinct responses to a given anti-tumor regime across different solid tumors. For instance, the immune-desert TMEs have been clinically observed to be non-responsive to neo-adjuvant immunotherapy as opposed to the immune-rich TMEs [9, 10].

Although a rigorous evaluation of the fibro-rich TME response to immunotherapy is scarce in the literature, several studies have identified the abundance of cancer-associated fibroblasts in non-responsive (to immune therapy) TMEs [11–14]. Despite the clinical observation of the TME subtypes, the exact genetic or epigenetic mechanisms behind a specific compositional environment remain unclear. Further, the exact mechanisms that explain the relation between a given TME subtype across the solid tumors and the responsivity of neo-adjuvant immunotherapy can improve the understanding of the disease and aid in synthesizing TME subtype-specific combination strategies.

The existing model-based approaches for addressing the aforementioned questions mostly focused on a specific part of TME [15–19]. The cell-level representation of the model assumes functional homogeneity across a given cell type which may not be reliable given the fact that two different cellular states of a given cell type can perform opposite functionalities [8, 13, 20–22]. Further, the existing modeling efforts do not provide a mechanistic explanation behind the emergence of different, apparently static, TME subtypes that, in turn, govern the response to immune therapy. This necessitates a conceptual framework that can (a) map the phenotypes relating to solid tumors to relevant modeling concepts and (b) be generalized to a wide range of bio-modeling rules.

Motivated by the limitations and the scope of the literature, this work operates at the cell state level, wherein we reconstruct a minimal interaction atlas common to the majority of solid tumors from the literature. Considering the populations and concentrations of the different cellular states and the associated molecular species, respectively, as the states, we construct the symbolic state space representation of the network. Further, (a) mapping the phenotypic variability of the TME to different properties of the underlying dynamical system and the subsequent (b) translation of these systems-guided conditions to structural underpinnings using the wealth of algebraic graph theory unravels the novel mechanisms behind the structural modulations crucial for the TME diversity, and the emergence of primary and acquired resistance. Finally, the systems theoretic treatment of the problem enables us to propose novel combination therapies that may improve the clinical outcome compared to sole ICI therapy. To this extent, we do not impose any particular kinetics on the dynamical system, barring some minimal assumptions. However, to validate the veracity of the theoretical predictions, we use an approximate cell-state transition-based model for the entire TME for solid tumors.

## 2 Results

Given the descriptive information in the literature about the constitutive units and their possible interactions, we propose the overall interaction atlas common to a wide range of solid tumors in Fig. 1 [1,6,11,13,20,23–74]. As discussed earlier, the cell state abstraction enables us to work with the level of granularity sufficient to characterize each unit’s role inside the TME precisely. Additionally, the cell state-level framework reduces the dimensionality of the problem, rendering it amicable for a generic, theoretical analysis [75]. We then propose a formalism that connects the relavant properties of the solid tumors to the important concepts of systems theory (Fig. 1).

**Fig 1.**
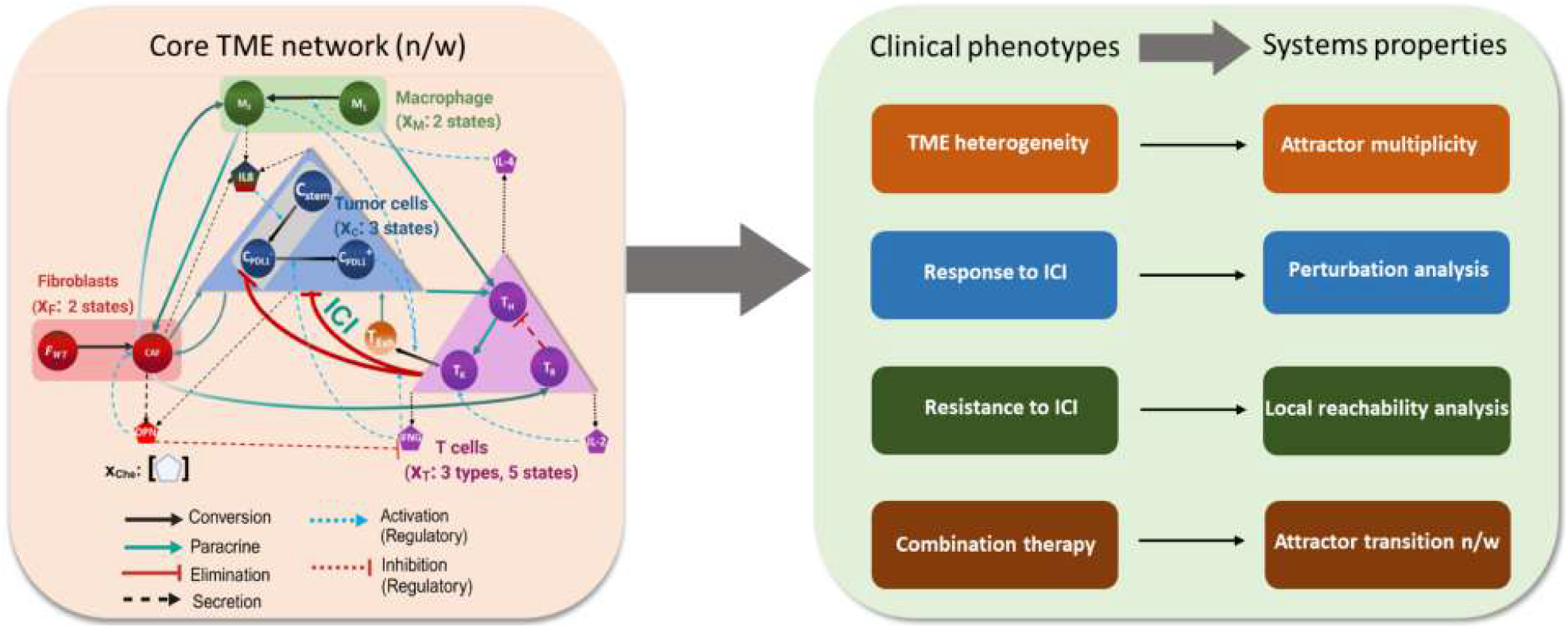
Workflow of the proposed formalism. We begin with the reconstruction of the cell-state-specific TME network for solid tumors. The nodes are either the cell states or the molecular species, whereas the edges represent diverse forms of interactions. The acronyms C_0, C_PDL-, and C_PDL+ denote the stem, PDL1^−^, and PDL1^+^ tumor cell states. T_K+, T_K-, T_Help, T_Reg, and T_Ex refer to PD1^+^ killer, PD1^−^ killer, helper, regulatory, and exhausted T cells, respectively. CAF, F_WT represents the pro-invasive cancer-associated fibroblasts and wild-type fibroblasts, respectively. The circles M_1 and M_2 refer to M1 and M2 phase macrophages, respectively. IL-2, IL-8, LIF, IRF8, OPN, Lac, ICAM1, and IFNG stand for interleukin 2, interleukin8, leukemia inhibitory factor, interferon regulating factor 8, osteopontin, lactate, intercellular adhesion molecule-1, and interferon-gamma respectively. Given the network structure, we set out to map the defining features of the solid tumor to pertinent concept of systems therory.

### 2.1 Characterizing the attractor landscapes: TME subtypes and its variants

As mentioned previously, different compositional possibilities within the TME are critical in determining the response to immunotherapeutic treatments [8]. Interestingly, despite the enormous compositional possibilities, a classification of the TME components across patients reveals the existence of a few distinct tumor microenvironments that have distinguishable effects on the ICI treatment [8, 14]. From a systems theoretic perspective, a finite number of TME phenotypes can be explained by conceiving the tumor microenvironment as a multistate dynamical system, wherein the attractors of the system represent the clinically observed TME compositions. Further, the ICI treatment outcomes for different TME subtypes are equivalent to the response to a specific perturbation analysis for the TME system around the neighborhood of different attractors of the pre-perturbed dynamical system.

With the proposed line of action and the given interaction map in Fig. 1, the underlying dynamical system can be represented as

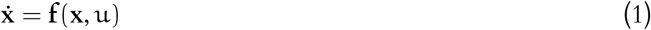

where **x** ∈ ℝ^N^ represents the states denoting the population and concentration of cellular states and the molecular species, respectively, and u(t) is the anti-PD1 drug concentration.

Considering the different interaction possibilities such as proliferation, paracrine, transition, elimination, and death, the above dynamical system given by (1) can be decomposed as

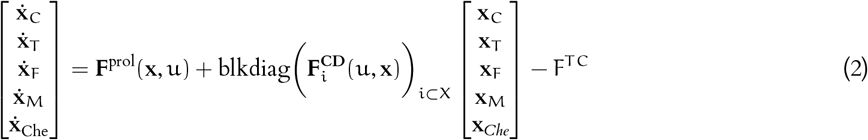

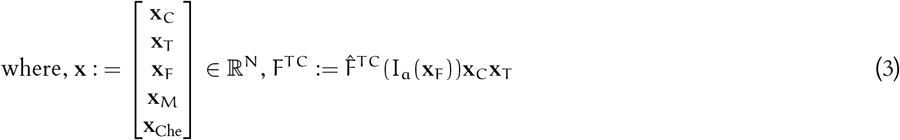

where, 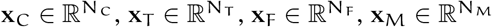 represent the population of the cell states pertaining to the tumor cells, T cells, fibroblasts, and macrophages respectively whereas 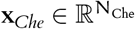 denotes the concentration of the molecular species. The function ‘blkdiag(.)’ returns a block diagonal matrix with the specific block matrices as diagonal elements representing the transition-like interactions in the TME. The set X contains all the node species. The function F^TC^ depicts the elimination action of the tumor cells by the killer T cells. Further, the quantity I_a_ refers to *immune accessibility index* that governs the effective availability of the cytotoxic T cells in the proximity of the tumor cells. Further, it is well known from the literature that immune accessibility is modulated by the cancer-associated fibroblasts (CAF) [14].

It is to be noted that the block diagonal structure imposes the fact that transition between two different cell states can happen within the same cell type. **F**prol(**x**) : **R**N → **R**N is the proliferation flux vector defined as

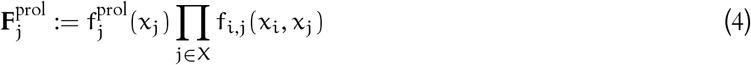

where, 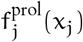 is the intrinsic resource-based proliferation rate for x_j_ and f_i,j_ is the flux modifier through paracrine and ligand-receptor based interactions between x_i_ and x_j_.

#### 2.1.1 Low proliferation, high exhaustion: Immune/Fibro desert

Given the local class 𝒦 (zero at origin, and locally monotone increasing with respect to x_j_) property of 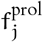, the dynamical system in (2) transforms into a *degenerate system*— There exists at least one steady state solution containing zero T cell population— We denote the set of such solutions 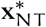. Similarly, the set of solutions corresponding to zero steady-state population for CAFs are denoted as 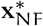 respectively. Intuitively, the sets 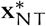 and 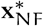 resemble the well-known immune desert and fibro desert TME subtypes. Further, this does not eliminate the non-zero steady-state solutions for the TME system.

It is to be noted that the mere existence of 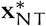, and 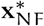 can not be directly linked with the clinical observation of immune and fibro desert TME for the additional stability conditions need to be met for any steady state to serve as a potential subtype. We show that under Assumptions 1-3, there exists a set of closed-form conditions constructed from the functional forms corresponding to the proliferation and conversion/death of killer T cells and fibroblasts that (Fig. 2) can serve as the *necessary condition* for 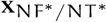 to contain at least one stable steady state. Importantly, these conditions have important biological implications. The necessary condition for immune desert-type attractors implies a critically low initial (when the killer T cell count is low) proliferation rate compared to the exhaustion and death rate of killer T cells. We term this scenario low-proliferation-induced immune desert. Further, our analysis also suggests that a threshold exhaustion rate exists for a given proliferation form with an arbitrarily high proliferation rate beyond which the tumor microenvironment transforms into an immune desert subtype— This can be distinguished as an exhaustiondriven immune desert.

**Fig 2.**
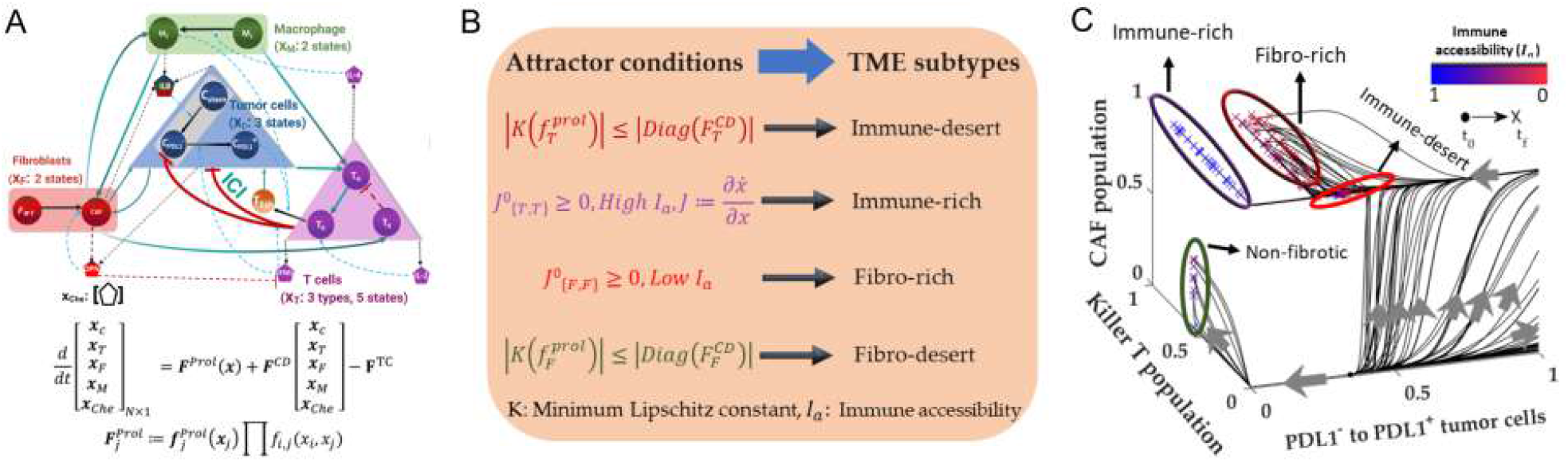
Multiple attractors explain TME subtype heterogeneity. Given the TME network in (A) we derive the closed-form conditions based on the generalized dynamics in (B) to find out the conditions for different attractor sets resembling clinically observed TME subtypes. (C) A computational illustration of different attractor possibilities with logistic proliferation and Michaelis-Menten type interaction rate. The fibro rich and immune rich attractors are distinguished by increasing immune accessibility. A low immune accessibility can also manifest as high PDL1-/PDL1+ tumor cell ratio

#### 2.1.2 Immune accessibility distinguishes immune-rich from fibro-rich

The proposed formalism establishes the viability of obtaining immune and fibro desert TMEs. Interestingly, for TMEs violating the desert conditions (denoted as non-fibro/immune desert TMEs) do not adopt the immune (fibro) desert as stable, steady states. Further, Assumptions 1-3 guarantee the existence (refer to Theorem 1 of the Appendix) of a positive invariant set (*i.e*., any trajectory starting within the set shall always remain in the set for all forward time) in the positive orthant of the state space with the planes **x**_F_ = 0 and **x**_T_ = 0 strictly repulsive. This implies the existence of at least one stable, steady state or a stable limit circle that contains at least one point with a non-zero fibro and immune cell population— This substantiates the existence of at least one fibro/immune-rich phenotype.

It is to be noted that both the CAF and Killer T cell components corresponding to an immune (fibro)-rich component can be non-zero. In this context, our formalism suggests that the immune accessibility index distinguishes the fibro-rich attractor from the immune-rich counterpart. According to the proposed condition, a fibro-rich attractor is identified with a low immune accessibility index due to the fact that an abundant CAF population, apart from aiding in the proliferation of the tumor cells, can alter the geometry of the TME, especially through construction of CAF-barriers around the tumor cells. On the contrary, the immune-rich attractor does not facilitate significant barrier formation activities of CAF. A computational validation of our analysis has been shown in Fig. 2 (C) wherein we considered a logistic proliferation growth with Michaelis-Meneten-like interaction kinetics.

Although the concept of immune accessibility explains the emergence of two attractors (fibro and immune rich) of similar composition but opposite effects (as we shall explore later in this work), the calculation of the same is not straightforward. To circumvent this limitation in the accurate identification of ‘rich’ phenotypes, we show that the ratio of the PDL1-to the PDL1+ tumor cells can serve as a good distinguishing factor between the immune and fibro-rich attractors in a pre-ICI setting. Our analysis suggests that under low immune accessibility, the major share of the total tumor cells remain *unreachable* to the killer T cells irrespective of the PDL1 signature, leading to a higher PDL1-/PDL1+ tumor cell. On the other hand, the immune-rich scenario, is characterized by the major section of the tumor cells being accessible to the killer T cells. Therefore, in a pre-ICI setting, the PD1+ killer T cells eliminate the PDL1-tumor cells providing a competitive advantage to the PDL1+ tumor cells— This results in a very low PDL1-tumor cell component compared to PDL1+ tumor cells in an immune-rich attractor.

#### 2.1.3 Can multiple attractors coexist?

The conditions provided for the existence of different clinically-observed TME subtypes shed light on the compositional characteristics of the different possible attractors or *stable TME compositions* for the dynamical system in (2). We here investigate the possibility of the coexistence of multiple attractors (strictly, attractors of different compositional characteristics vis-a-vis the CAF and immune cell population) for a given dynamics. Note this question is of central importance in designing the combination therapy, for if the dynamics allow multiple different attractors with differing ICI responses, it may be possible for a particular perturbation strategy such that the solution trajectory can be transitioned from an ICI-resistant attractor to a responsive attractor.

With an assumption of bounded kinetics, we propose a set of sufficient conditions for the given dynamical system in (2) such that the solution trajectories always converge on to an immune desert attractor for all non-negative initial conditions– We call this as absolute immune desert attractor. A critically low net proliferation flux of the killer T cells ensures the existence of the absolute immune desert. Evidently, the condition of the absolute immune desert is a stronger condition than the necessary condition for the existence of at least one immune desert attractor.

Conversely, we also lay out a set of sufficient conditions that ensure the co-existence of an immune-rich attractor (non-zero killer T cell component) along with an immune-desert attractor. In this scenario, the final state of the solution to (2) is conditioned on the initial condition. In this scenario, we term the immune-desert attractor as the conditional immune-desert attractor.

The necessary conditions provided for the absolute immune/fibro-rich attractors imply the zero killer T cells/ CAF can not be in a stable, steady state– This eliminates the co-existence of a desert-like attractor. Therefore, we here ask a very specific question of whether it is possible to have multiple attractors with the following immune/fibro-rich compositional characteristics i) Low fibrotic-low immune cells, high fibrotic-high immune cells and ii) low fibrotic-high immune cells, high fibrotic-low immune cells.

Our analysis suggests that under an aggressive immune activity, both scenarios are not possible (Claim 1 and Remark 2 of the Appendix) for the dynamics under Assumptions 1-3. However, the immune accessibility index (I_a_) modulates the steady-state population of the tumor cells. To demonstrate the analytical conclusions, we also device a simulation study in Fig. 3 (D-F) of five hundred different initial conditions with the logistic growth and Michael-Menten like kinetics. We ensure the proposed conditions for absolute and conditional existences in all the scenario. To summarize, the systems-theoretic investigation of the proposed network at the pre-ICI stage reveals five effective TME subtypes: i) desert, ii) fibro desert, iii) immune desert, iv) fibro rich and v) immune rich. Further, depending on the dynamical forms and the parameters, these attractors can be global or co-exist within the state space.

**Fig 3.**
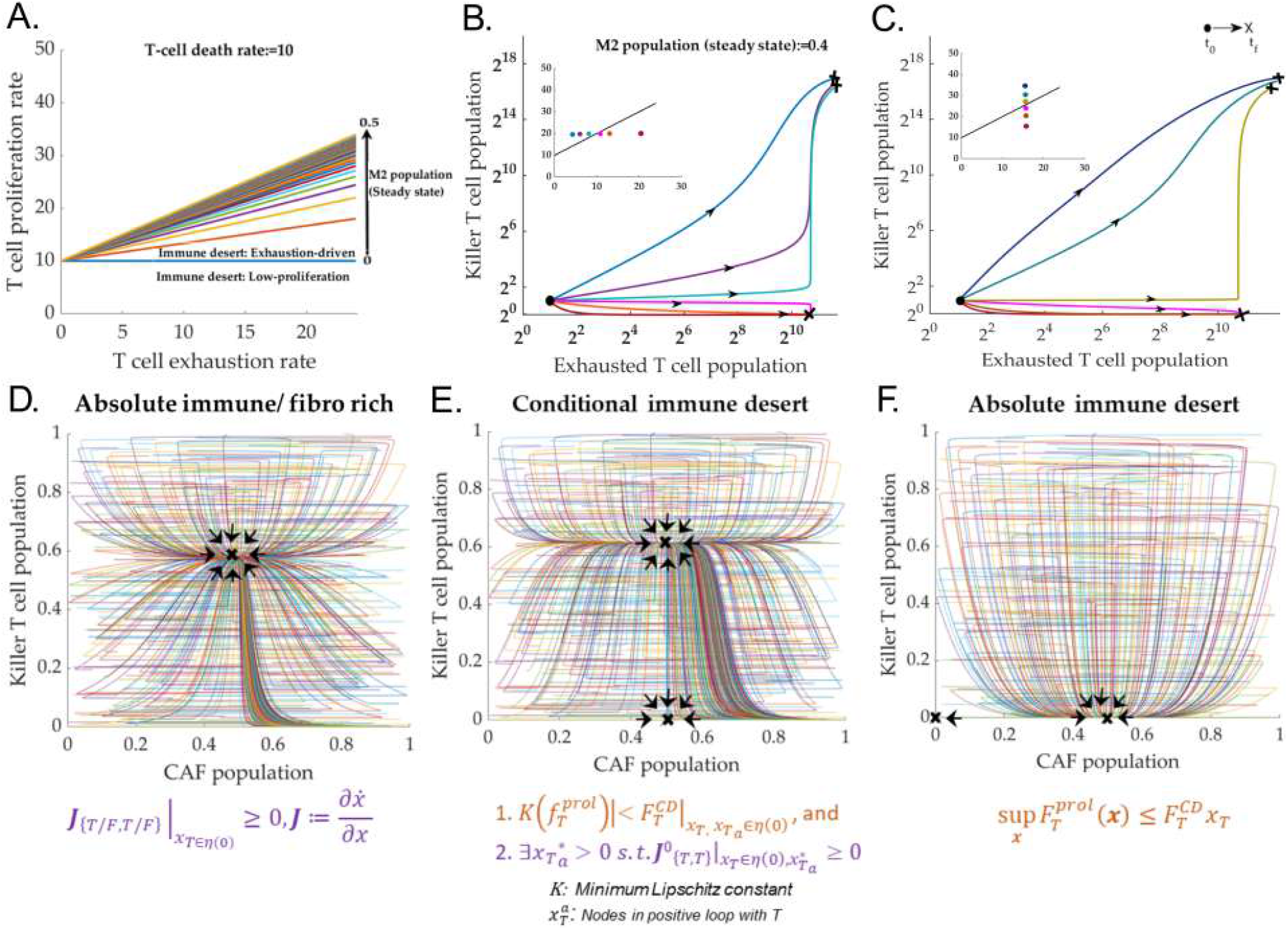
Diversity with and co-existence of attractors. (A-C) Low-proliferation and exhaustion-driven immune desert: Existence of an M-2 dependent threshold exhaustion rate given any proliferation rate for immune depletion. (D-F) Demostration of attractors with different compositional characteristics co-existing. We also lay out the underlying mathematical conditions for the possibility of co-existence.

### 2.2 Anti-PD1 perturbation: Reachability governs response, reduction, and no response

According to the proposed model, the anti-PD1 drug facilitates the conversion to PD1^−^, killer T cells from its PD1+ counterpart. The elimination of the PD1 program enables the killer T cells to destroy the tumor cells irrespective of the PDL1 expression.

Our analysis suggests the binary question of whether a particular TME subtype is responsive to anti-PD1 therapy can be resolved by performing a reachability analysis of the system around the pre-ICI attractor. From a graph-theoretic perspective, the reachability analysis around the attractor determines if there is at least one path from the anti-PD1 dosage to all the tumor cell states in the influence graph constituted from the Jacobian of the right-hand side in (2) with respect to all the states. In a strict sense, the mere existence of a path from the anti-PD1 to tumor cell states is not enough, for it does not say anything about the effective sign of the paths. The sign of the forward path dictates the overall effect of anti-PD1 on the tumor cell states. For instance, if the net path sign is positive, the tumor cell states will likely increase post-ICI therapy. Therefore, along with the reachability condition, we propose that the cumulative path sign from anti-PD1 to tumor cell states has to be negative in the digraph constituted from the Jacobian analysis. From algebraic graph theory, the minor of the component in the Jacobian matrix concerning the direct influence of tumor cell states on the anti-PD1 provides the effective sign information of all the forward paths from anti-PD1 to tumor cell states.

#### 2.2.1 Immune desert: No reachability, no response

We first begin with the immune desert subtype, which, according to our analysis (Proposition 1 Appendix), is viable when the death and exhaustion rate of the immune cells is higher than a critical value (Lipschitz constant) associated with the net proliferation rate. Our analysis suggests that the attractors with immune desert composition render the localized influence graph (digraph induced by the Jaboian matrix evaluated around the attractor) with no forward paths from the anti-PD1 to the tumor cell states. This is primarily due to the non-existence of any link from the anti-PD1 to all the killer T cells owing to the absence (or very few) of PD1+ killer T cells in the neighborhood of an immune desert attractor.

#### 2.2.2 ICI may facilitate complete recovery in fibro-desert region

The fibro desert phenotype, on the other hand, consists of the Killer T cells, Tumor cells, and the killer T cells. The TME is also depleted of inflammatory cytokines such as OPN, IL-8, and LIF due to the absence of CAF and the shortage of M2 macrophages due to the lack of major growth-promoting interactions for M2 macrophages, such as the CAF-TAM interaction [22]. Applying the reachability analysis and the subsequent path sign calculation, we obtain that the overall sign of the path from anti-PD1 to all the tumor cell states is strictly negative (Fig. 4 (A)). Therefore, we conclude that fibro-desert constitutes a favorable scenario for anti-PD1 therapy. However, whether the post-ICI tumor cell population following an ICI therapy goes to zero is governed by the residual resource concentration. In this context, we define the high immune activity region for a given level of T cell-driven clearance rate as the set of resources bounded by the maximum amount of resources till the immune-accessible tumor cell population converges to zero. Further, immune-accessible tumor cell states are defined as the tumor cell states that are (a) reachable from the anti-PD1 and (b) the effective path sign from the anti-PD1 to the tumor cell state is negative. Interestingly, our analysis suggests that the application of anti-PD1 significantly extends the high immune activity region, implying the possibility of complete recovery post-ICI therapy for a wide range of resource concentrations. The simulation study Fig. 4 (B-C) with logistic growth and rational interaction kinetics agree with the theoretical findings.

**Fig 4.**
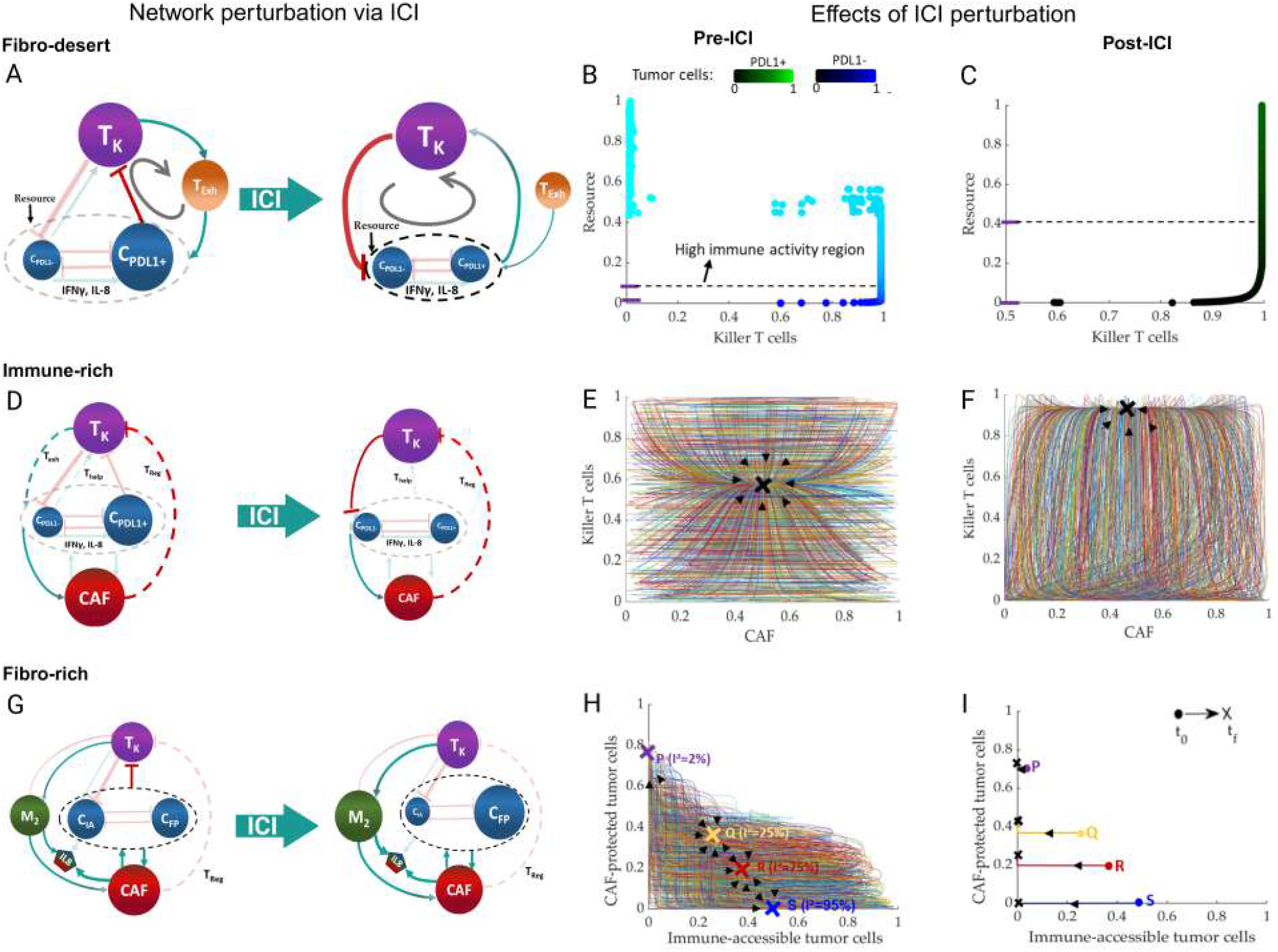
Response to ICI: Perturation around the pre-ICI attractor. (A) Influence graph around a fibro-desert attractor. The graph obtained from the computing the Jacobian of the dynamics with with respect to the states and evaluating at the fibro-desert attractor. The green and red arrows mean positive and negative influences respectively. (B-C) The high immune acticity region increases post-ICI for fibro-desert attractor. (D-F) The anti-PD1 reprograms the influence graph of immune rich attractor towards a reduction of both the tumor and CAF cells (G-I) The effective path from the anti-PD1 to the tumor cell states in the influence graph for fibro-rich attractor are almost zero and positive thereby indicating towards almost no response and an increase in CAF-protected tumor cells.

#### 2.2.3 Immune to fibro rich: Changing response spectrum

As established before, the ‘rich-’TME phenotypes can be distinguished by the immune accessibility index. The immune accessibility to the tumor cells is governed by the barrier formation rate of the resident CAF population. A lower barrier formation rate, implying higher immune accessibility, results in an attractor with immune-rich properties *i.e*., zero or very low PDL1-/PDL1+ tumor cell ratio. Our analysis suggests that although the killer T cells eliminate the PDL1-tumor cells in the pre-ICI setting, the effective path sign for the killer T cells to the CAF is non-negative primarily due to the pro-tumor activity of the exhausted T cells. However, an anti-PD1 intervention, eliminates the T cell exhaustion program and establishes a negative path from the killer T cells to CAF. Therefore, we conclude that the ICI intervention results in a decrease of CAF population along with the tumor cells when compared to the pre-ICI counterpart. Fig. 4 (D-F) provides computational illustration of the ICI perturbation of immune rich attractor. Interestingly, our analysis suggests that under certain closed-form conditions on the proliferation rate of the killer T cell, the ICI-based intervention cannot fully remove the tumor cells even in the presence of high killer T cell activity. Remark 5 of the appendix elucidates the scenario in a comprehensive manner.

On the other hand, the reachability analysis around a fibro-rich attractor reveals that although the net sign of the path from the anti-PD1 species to the immune-accessible tumor cells is strictly negative, the path to the CAF-protected tumor cells is of non-negative sign— This is primarily because (a) the absence of killer T cell-driven elimination of CAF-protected tumor cells and (b) the net path from killer T cell to immune accessible tumor cells to CAF protected tumor cells is positive owing to the resource and space competition between the tumor cells. Therefore, in the fibro-rich scenario, the post-ICI immune-accessible tumor cells decrease at the cost of an increase in the population of CAF-protected tumor cells. At extremely low immune accessibility levels, the post-ICI total tumor cell population remains almost identical to its pre-ICI counterpart. Fig. 4 (G-I) supports the analytical findings.

### 2.3 Overcoming non-response: Leveraging structural theory for attractor transition network

As discussed in the previous sections, the reachability analysis along with the examination of the path (from anti-PD1 to tumor cell states) reveals that while the attractors resembling fibro-desert and immune-rich TME subtypes respond to ICI the immune desert and fibro rich subtypes remain largely insensitive to ICI theray.

Therefore, designing approapriate theraputics to transition any trajectory towards the fibro-desert or immune-rich attractors costitute the potential for combination therapy. From the perspective of control theory, this problem is equivalent to the design of appropriate control that can drive any trajectory to the desired attractor. Fiedler *et al*. (2013) proved that under certain bounded dynamics with non zero decay condition, the node set constructed by considering at least one node from each feedback loop (termed as the feedback vortex set) in the network serve as the sufficient set of node overrides (in the absence of source nodes) for driving the solution trajecotries to the desired attractors. We show the assumptions made in this work constitutes a suitable premise for application of the aforementioned result.

Since overridding the tumor cells, CAF and killer T cells in a direct fashion is out of the scope of this work we work with a subset of the feedback vortes set. We chose the set 𝒮^FVS^ = {OPN, IL − 2, M_2_}.

Interestingly, our analysis suggests that under some closed-form conditions on the dynamics of CAF (Proposition 6 of the Appendix), there exists a threshold OPN elimination rate for which the fibro-rich TME sub-type transforms into a fibro-desert subtype. Further, the depletion of CAFs leads to an increase in immune accessibility. Therefore, for a non-immune-desert scenario, the OPN removal modifies the TME into an immune-rich TME phenotype (Fig. 5 (C-D)).

**Fig 5.**
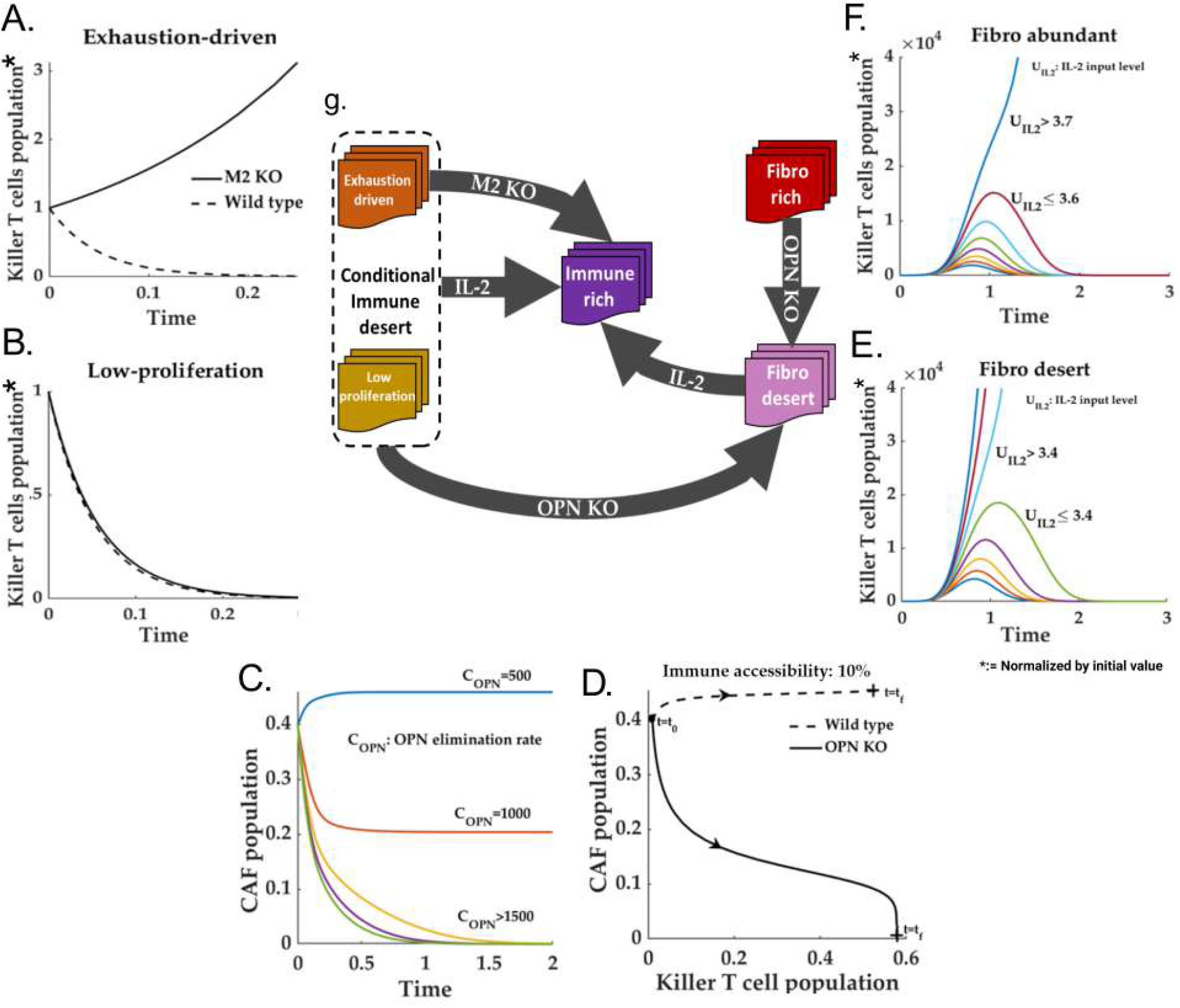
Study of possible transitions. (A-B). An M2 knockout can drive an exhaustion-driven immune-desert to an immune-non-desert phenotype if the killer T cell proliferation is beyond a critical level. (C-D) OPN deletion results in a fibro-desert-like TME, even in a TME with very low immune accessibility. (E-F) IL-2-based intervention can result in the transition from any immune-desert attractor to an immune-non-desert attractor. Further, the minimum IL-2 required for this transition is reduced in a fibro-desert situation.

Further, we also claim that an M2 macrophage knockout can revive all the solution trajectories out of exhaustion-drive immune desert attractor as long as the initial slope (with respect to the population) of proliferation is greater than the death rate of the killer T cells (Fig. 5 (A-B)).

On the other hand, under certain conditions on the proliferation rate of the Killer T cells, we show that there exists a one time IL-2 insertion rate (Proposition 4 of the Appendix) that can destabilize the immune-desert subtype. However, a constant injection of IL-2 is not realistically possible for the well-known toxicities of IL-2. Therefore, we exploit the reinforcing loop between the killer T cell and IL2 to propose a constant one time IL-2 insertion amount (Proposition 5 and Lemma 7 of the Appendix) for a given dynamics that can drive the state trajectories out of the domain of attraction of the immune-desert attractors (Fig. 5 (A-B)).

## 3 Discussion

A mechanistic understanding of complex diseases requires a conceptual framework that can map the defining features of the disease to the language of dynamical systems theory. The systems theoretic tools aid in engineering possible therapeutic strategies for further experimental validation. In this work, we provided a system-theoretic framework that translates the pertinent properties of the solid tumors such as TME diversity, prognostic heterogeneity to the language of systems theory which in turn, aid not only in possible explanations of the mapping between the TME subtypes to ICI response but also proposes combinatiorial therapeutic strategies to circumvent the emergence of ICI resistance.

For this purpose, we reconstructed the TME from the existing literature on single-cell analysis of solid tumor patients. By design, the cell-state-based formalism compresses the epigenetic information in a few dimensions and, at the same time, can capture a wide range of phenomena compared to its cell-level counterpart.

Considering the population and concentration of different cell states and molecular species as the states, we constitute the dynamical system associated with the core TME network.

Given the networked system,, we show that the apparent heterogeneity in the TME subtypes can be explained through different compositional possibilities of systems attractors in a pre-ICI setting. Thus, the specific conditions for the existence of a given attractor can provide a mechanistic understanding behind the emergence of the associated TME sutype. For instance, we show that for any given proliferation form of the killer T cells, there exists a critical exhaustion rate beyond which the TME settles to an immune desert configuration.

Given the potential attractor set, we investigated the possibility of co-existence of different attractors for a given set of dynamical forms and associated parameters— this translates to the question of whether a given epigenetic environment can lead to multiple outcome vis-a-vis the TME subtypes. We found that under certain conditions, the immune/fibro rich attractors can indeed co-exist with immune/fibro desert attractors. This result would be central in designing the subtype-specific combination therapies.

While the compositional diversity of the solid tumor TME can be mapped to the attractors of the autonomous system the response to ICI therapy amounts the perturbation analysis of the system around the pre-ICI attractor. We proposed the path-informed reachability-based analysis can be an effective tool for determining whether a given TME subtype is ICI resistant. Using this we were able to explain the well-known ICI resistance exhibited by the immune desert and fibro-rich subtypes. On the other hand, the Since, the co-existence of different attractors are possible, we used the structural theory by Feidler *et al*. (2013) to construct the feedback vortex set corresponding to the network. Based on the consideration of this work, we chose three nodes OPN, IL-2 and M-2 macrophage for perturbations. We show, under certain conditions involving the conditional existence of fibro-desert attractor, there exists a threshold OPN elimination rate beyond which all the solution trajectories near to the ICI resistant fibro-rich attractor are driven towards ICI responsive fibro-desert attractor. Similarly, an M-2 knockout can drive the system trajectories out of the exhaustion-driven immune desert to an immune-rich attractor. On the other hand, a one time IL-2 intervention can also break the deadlock of an immune desert (both low proliferation or exhaustion-driven) to steer the trajctories towards an immune rich phenotype.

The proposed formalism does not assume any particular rate kinetics or modeling rule and therefore can be generalizable to a wide range of biologically sensible modeling rues. Although the proposed formalism captures a wide array of experimentally observed phenomena, it does not explain some of the most crucial aspects of solid tumors, such as staging and metastasis. Therefore, the development of a comprehensive formalism incorporating intra-tumoral heterogeneity and metastasis can be a tempting area of future study.

## Appendix: From

### 1 Dynamics

The generalized representation of the population (concentration) dynamics for different cellular states and molecular species inside the TME can be formulated as

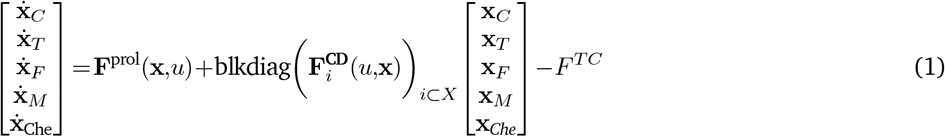

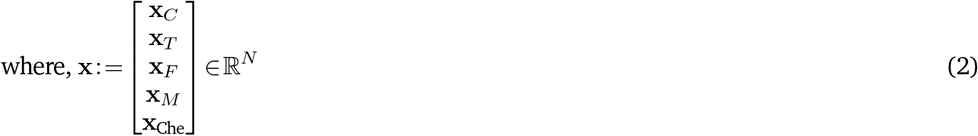

With

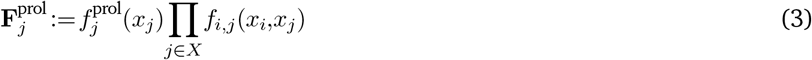

#### Assumptions

With the mentioned dynamics, we list out all the important assumptions we made at some point in this work.

1. Well-posedness: The flow associated with (1) satisfies the semi-group property, and the vector field is Lipschitz continuous concerning the states, and the input (*u*). This ensures the fact that the dynamical system in (1) constitutes a *well-po*s*ed* dynamical system [1].
2. Semi concavity: The proliferation flux 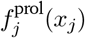 is semi-concave and class 𝒦 around the origin.
3. High killer T cell activity: Tumor cells susceptible to cytotoxic T cells converge to zero asymptotically.
4. Boundedness: The functions 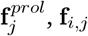 are bounded above.

### 2 Well-posedness and positive invariance of the TME system

The well-posedness of a dynamic system refers to the existence of a trajectory from the initial condition. Further, given an initial condition, the uniqueness of a given trajectory can be guaranteed as long as the flux is locally Lipschitz. Given the dynamics in (1), since the function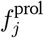 is concave and hence Lipschitz, the functions *f*_*i,j*_ are bounded to the function 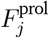 is also Lipschitz. Further, the additive terms 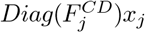 is linear. Therefore, the overall dynamic system is globally Lipschitz. Therefore, all the solutions from all possible compact sets in *t* exist, and for a given initial condition, the solution to the dynamical system is unique.

Positive invariance is an extremely important concept in the analysis of dynamical systems. A set *S* ∈ ℝ is said to be positively invariant for a dynamical system ⅅ if for initial conditions **x**_0_ ∈*S*, the solution **x**(*t*):= ⅅ (**x**_0_,*t*)∈*S* ∀*t*≥0

Further, the following result concerns the existence of positive invariant sets for the aforementioned dynamical system with the given assumptions.

#### Theorem 1.

*Under assumptions 1-4, with strictly positive death/ degradation rates, the dynamical system in (1) corresponding to the TME network in Fig. 1 admits at least one positive invariant set with origin as the boundary*.

Before proving the theorem, we prove the following dependencies.

#### Lemma 2.

*For two scalar variable x* ∈ ℝ ^+^ ∪{0} *and y* ∈ℝ ^+^ ∪{0} *with the following dynamics*

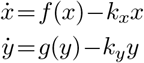

*where g*(*p*)≥*f*(*p*) ∀*p* ≥0 *with g*(0)=*f*(0)=0 *and k*_*y*_ *<k*_*x*_, *the following inequality holds x*(*t*)≤*y*(*t*) ∀*t*≥0 *if y*(0)≥*x*(0)≥0

*Proof*. We prove this using two steps. In the first step, let us define a variable *x*_1_ with the dynamics

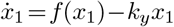

Further, let us define the difference variable *e*_1_ :=*x*_1_−*x*. The corresponding dynamics of *e*_1_ can be written as

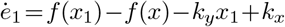

Further, at *e*_1_ =0 ⇒ *x*_1_ =*x*

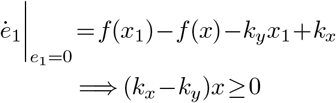

Therefore, the quantity *e*_1_ is always non-negative implying *x*_1_(*t*)≥*x*(*t*).

In the second step, we define another variable *e*_2_ :=*y*−*x*_1_. Therefore, the dynamics of *e*_2_ can be expressed as

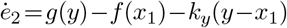

Further, at *e*_2_ =0 ⇒ *y* =*x*_1_

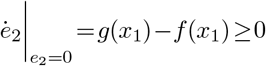

Therefore, ∀ *t*≥0, *e*_2_ ≥0 ⇒ *y*(*t*)≥*x*_1_(*t*)≥*x*(*t*). This concludes the proof.

#### Lemma 3.

*For a network with dynamics*

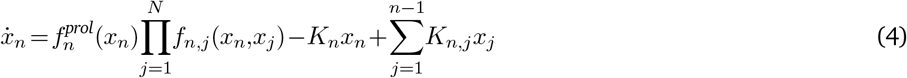

*with K*_*i,j*_ ≥0 ∀(*i,j*)=1(*i*)*N and K*_*i*_ *>*0. *The network consists of at least one positive invariant set if Assumptions 1-4 are satisfied.*

*Proof. For the variable* x_1_ *the dynamics can be written as*

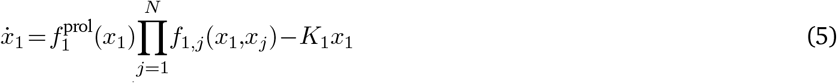

Let us define a set 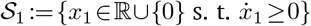. Since 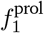 is locally class 𝒦 around the origin, the system is degenerate. Further, since 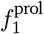 and *f*_1,*j*_ are bounded above with *B*_1_ and *B*_1,*j*_ respectively. From the Archimedean property of ℝ, there exists a 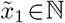 such that 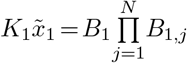. Further, since, 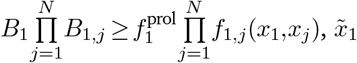 is an upper bound of 𝒮_1_. Therefore sup *S* exists. We denote it by 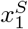. Therefore, *x*_1_ is always bounded by 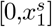.

Further, for *x*_2_, the set 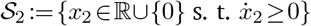 is bounded above by 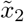 that satisfies the equation 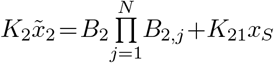 where, *B*_2_, and *B*_2,*j*_ are the upper bounds of 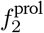 and *f*_2,*j*_(*x*_2_,*x*_*j*_). Therefore, the supremum exists and is denoted by 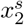.

Let us assume that the supremum exists for first *m* (2≤*m<N*) variables. Then, using the similar logic, the supremum also exists for the *m*+1 variables which can be obtained by solving the equation 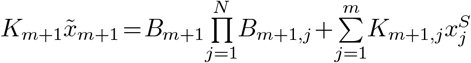.. where *Bm*+1, *B*_*m*+1,*j*_, and 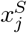 are the upper bounds of 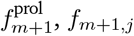,and the supremum value for *x*_*j*_ *j* =1(*i*)*m*. Therefore, by the theorem of mathematical induction, we prove that the hypercube 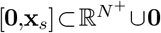 is a positive invariant set with respect to the dynamics. This concludes the proof.

#### Lemma 4.

*For two variables x*_1_ *and x*_2_ ∈ ℝ^+^ ∪{0}, *with the following dynamics*

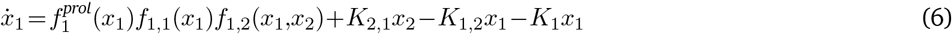

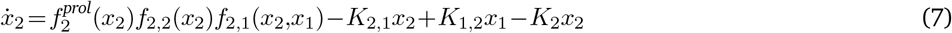

*There exists a positive invariant set if the dynamical system satisfy Assumptions 1-4*.

*Proof*. Given the dynamics, the variable (*x*_1_+*x*_2_) obeys the following dynamics

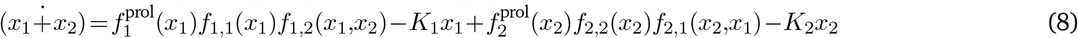

Now, since 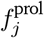 is concave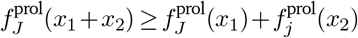.Since, *f*_*i,j*_ is bounded above by *B*_*i,j*_ according to Lemma 2 the variable *C* following the dynamics

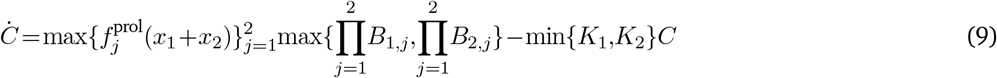

follows the relation *C*(*t*) ≥ *x*_1_(*t*)+*x*_2_(*t*) ∀*t* ≥ 0 and *C*(0) ≥ *x*_1_(0)+*x*_2_(0). Now, from claim 2, *C* is bounded above by the quantity obtained from solving the equation 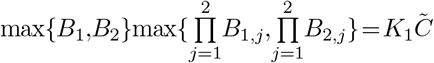.Therefore, the quantity *x*_1_(*t*)+*x*_2_(*t*) is also bounded by 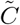. Further, the class 𝒦 nature of 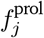 around the origin ensures the semi-positive nature of *x*_1_ and *x*_2_. Combining these two, the compact set 0≤(*x*_1_+*x*_2_)≤*C* with *x*_1_,*x*_2_ ≥0 is a positive invariant set for the above dynamics. This concludes the proof.

#### Lemma 5.

*Suppose* **x**:=[*x*_1_,*x*_2_,…,*x*_*N*_ ]^*T*^ *with x*_*i*_ ∈ ℝ ^+^ ∪{0} *follows the dynamics*

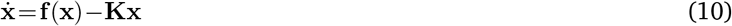

*where*, 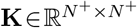 *is a diagonal matrix and* **f** (**x**)=0 *at x* =0. *Further, let* **p** ∈ ℝ ^*P*^ *is another set of variables with arbitrary dynamics. Assume the dynamics of* **x** *contains a positive invariant set* 𝒮_*x*_ *containing the origin. Then, the dynamical system for a new set of variables* 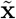

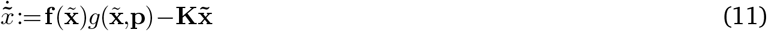

*contains at least one positive invariant set if* **g**_*j*_ *is bounded above*

*Proof*. Given **x**_*i*_ are bounded above then there exists a 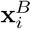 such that 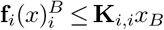. For each **x**_*i*_, define another variable 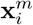 with the dynamics 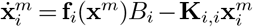 where, *B*_*i*_ is the upper bound for **g**_*i*_. Since *B*_*i*_ *<* ∞ and sup(*S*_*i*_) exists in ℝ where *S*_*i*_ := {*x*_*i*_ ∈ ℝ ^+^ such that **f**_*i*_(**x**) ≥ **K**_*i,i*_**x**_*i*_, the supremum of the set 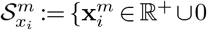, such that 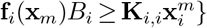 also exists.

Further, since, **f**_*i*_(**y**)*B*_*i*_ ≤ **f**_*i*_(**y**)**g**_*i*_(**y**,**z**) ∀**y** ∈ ℝ ^*N*^ ∪ **0**, according to Lemma 2 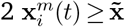. Therefore, 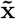 is also bounded above. This concludes the proof.

With these four claims, the proof for Theorem 1 is immediate

*Proof*. Given the dynamics in (1), Assumptions 1-4, and the TME network in Fig. 1 of the main manuscript, the tumor cell module contains three states— stem, PDL1^+^, and PDL1^−^ with the stem being transitioned to PDL1^+^ and PDL1^−^ tumor cells, and PDL1^−^ tumor cells transition into PDL1^+^ tumor cells. Therefore, using the result from Lemma 3, we conclude the tumor cell state population is always bounded above. Similarly, in the T cell module, the killer T cells transition into exhausted T cells. All the other T cellular states in T do not engage in transition. Therefore, using Lemma 3, we conclude that the T cell population is also bounded above. Further, the macrophages consist of transitions between both M1 and M2 cell states. The scenario for fibroblasts with two cell states, wild type and pro-invasive CAFs, is similar. This follows the dynamics as laid out in Lemma 4. Therefore, it is always bounded above. Combining all these and Lemma 5 to cover the communications between different modules, we conclude that the entire TME system with dynamics laid out in (1) contains at least one positive invariant set. This concludes the proof.

### 3 TME 𝒮ubtypes

#### Proposition 1.

*Under assumption 1,2, if the set* 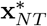 *contains at least one stable, steady state for the dynamical system in Eq.2 of the main manuscript, then there exists a functional f*_1_ *such that*

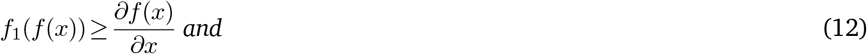

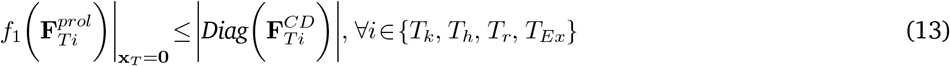

*Proof*. Since 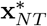 does not contain any T cells, the Jacobian 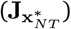 of the dynamical system in (1) of the main manuscript evaluated around 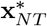 can be expressed as

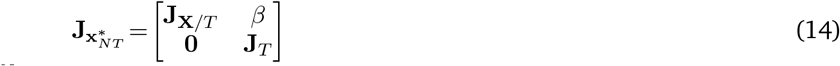

where, 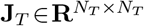 and 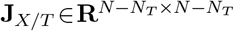, are the Jacobian of 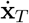 with respect to **x**_*T*_ and Jacobian of the dynamics of all the nodes (*X/T*) except the T cells with respect to **x***/***x** evaluated at 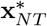, respectively. Due to the upper triangular nature of 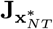, we can write the following

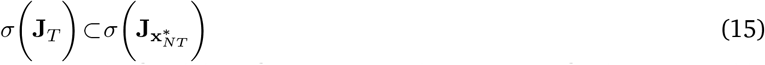

where, *σ* denotes the spectrum of a square matrix. From (15), the feasibility of 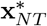 vis-a-vis the existence of at least one stable equilibrium is conditioned on the Hurwitz property of **J**_*T*_.

Further, it can also be shown with a simple calculation t t the matrix **J**_*T*_ is diagonal, with the diagonal element being

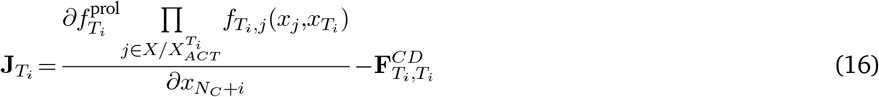

where the set *X*_*ACT*_ ⊂*X* is the set containing the nodes that facilitate the proliferation of *i*^th^ T cell species via autocrine interactions. Owing to the autocrine interaction, around the steady state of 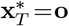, the concentration of all the species in *X*_*ACT*_ is zero.

For stability around the origin, there exists an ope neighborhood of the origin 𝒮_o_ such that the following is true

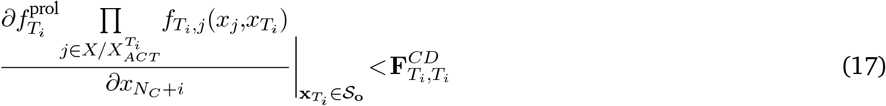

According to assumption 1, *f*^*prol*^ is semi-concave, which ensures Lipschitz continuity. Further, the function *f*_*i,j*_(*x*_*i*_,*x*_*j*_) is Lipschitz. Therefore, the combination 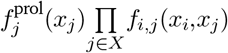 is also Lipschitz. Further, since, 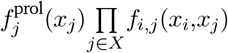 is global Lipschitz it is locally Lipschitz.

We define, the function *f*_1_(*x*) as

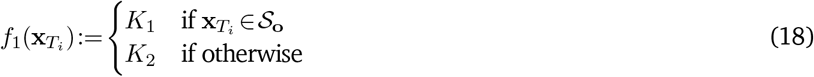

where, 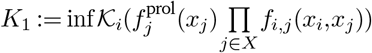 in the interval 𝒮o, 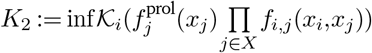 in ℝ ℝ ^+^*/*𝒮_o_. 𝒦_*i*_ are the local Lipschitz constants. By definition of Lipschitz’s constant 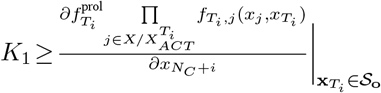 and 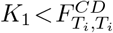 (*K*_1_ is the infimum).

This completes the proof.

#### Corollary 1.

*For the network dynamics in (1) obeying Assumptions 1-4, and strictly positive death/ degradation rates, there exists at least one stable attractor or limit circle with zero killer T cell population if the following condition is satisfied*

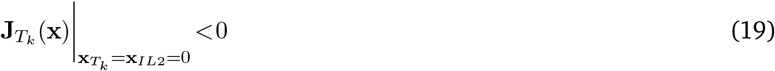

*where 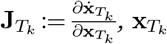, and **x** _IL2_refers to the population of killer T cells and concentration of IL-2 respectively*.

*Proof*. From the virtue of the dynamical system in (1) being a degenerate system, 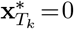 is a steady state for the T cell modules. Since, according to the core TME network IL-2 is secreted exclusively from the killer T cells 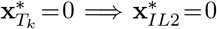. Since, the (19) is assumed, the quantity 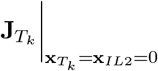 is always negative throughout the plane. Therefore, the plane 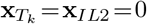 is positive invariant. Further, the assumption of strictly positive death/ degradation rates of all the network components satisfies the premise of Theorem 1. Therefore, there exists a positive invariant set even in the plane 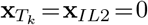. Therefore, the overall dynamical system contains at least one attractor or a stable limit circle in the plane 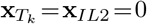. This concludes the proof.

#### Remark 1.

*Proposition 1 provides a closed-form condition for the stability of immune/fibro desert. Nevertheless, it does not eliminate the possibility of a non-desert-like attractor in the state space. The condition used in* (13) *takes care of the local stability: there exists an open neighborhood of **x**_T_ such that all the solutions starting within that neighborhood converge to a steady state in 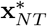. We call this open neighborhood as domain of attraction. However, the solutions sufficiently far from the domain of attraction of 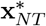 may not converge to a desert state. We shall exploit this aspect in the design of TME-specific perturbation strategies. Further, the condition for absolute desert can be obtained if the following condition is satisfied*

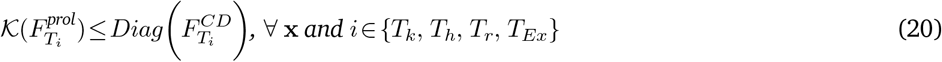

*where, 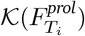 is the Lipschitz constant associated with 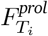. This guarantees that the vector field 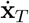 remains negative (barring the origin) throughout the state space. Therefore, the only stable, steady state is when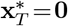. Note, (20) is satisfied by (13) for 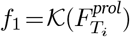, but the converse is not true*.

#### 3.1 Desert modalities

The following results shed light on different possible mechanisms for the cell state trajectories within a TME to converge to an immune-desert attractor. Before proceeding with the main mathematical result we propose the following definition

##### Definition 1.

*The TME network with the dynamics in* (1) *satisfies an M2 non-desert subtype if there exists a* 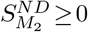 *such that the following condition is satisfied*

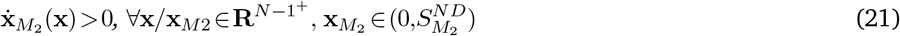

##### Lemma 6.

*Given all the parameters of a non-fibro-desert TME network shown in Fig. 2, the main manuscript with the associated dynamics (1) obeying Assumptions 1-4, there exists a threshold conversion rate, from the killer T cells to the exhausted T cells beyond which, under M2 non-desert scenario, the resultant dynamical system contains at least one attractor resembling immune desert TME phenotype*.

*Proof*. The dynamics of the killer T cells 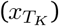 can be written as

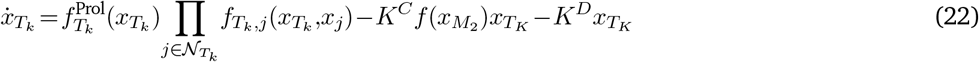

where, 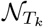 refer to the incoming neighbor nodes of the killer T cell. Since the TME subtype is non-fibro-desert, the point 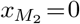 can not be figured in any system attractor. Therefore, any trajectory within the arbitrary neighborhood of 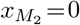 shall move away from the point. Given the state trajectory **x**(*t*) to the dynamical system in (1), we construct the set 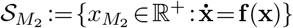 By construction, the set 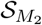 is bounded below by zero. Using Theorem 1, all the trajectories starting within the positive invariant set of the positive system in (1) have at least one point in the state space with a non-zero *M*_2_ component. Therefore, the set 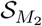 is non-empty. This implies 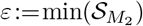 exists and is positive.

Therefore, for any *M*_2_ population 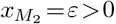, the term *f*(*ε*) is strictly non-zero positive and minima of *f* throughout the state space trajectory. Further, from the Archimedean property of ℝ, there exists an 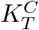 such that the following condition is satisfied

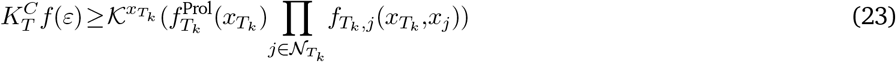

where, 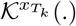 is the Lipschitz constant with respect to 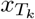. It is to be noted that (23) satisfies the desert condition laid out in Corollary 1. Therefore, any exhaustion rate beyond 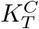 reprograms the TME into an immune-desert subtype. This concludes the proof.

### 4 Investigating the CAF-T landscape: Non-desert phenotypes

In this section, we aim to work out the non-desert conditions *i.e*., the existence of attractors and their properties. The following remark sheds light on the existence of non-desert attractors. We first start with a formal definition of non-desert attractors

#### Definition 2.

*An attractor set* 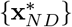 *is called non-desert attractor set to the dynamical system in* (1) *if the CAF and killer T cell components of* 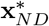 *are strictly non-zero*.

#### Remark 2.

*The proposed formalism establishes the viability of obtaining immune and fibro desert TMEs. Interestingly, for TMEs violating the desert condition in (denote as non-fibro/immune desert TMEs) may not adopt the immune (fibro) desert as stable, steady states. Using the same logic, the desert can also be ruled out as a viable TME composition. Further, Assumptions 1-4 guarantee the existence of a closed interval* [**0**,**x**_*max*_] *which is positive invariant (Theorem 1) with the plane* **x**_*F*_ =**x**_*T*_ =0 *strictly repulsive. This implies the existence of at least one stable, steady state or a stable limit circle that contains at least one point with a non-zero fibro and immune cell population*.

It is to be noted that the existence of an attractor, hitherto, is considered with respect to different parameters characterizing the dynamics. In contrast, we here investigate the possibility of multiple non-desert attractors in the state space with a given parameter set. In other words, we characterize the cardinality of the non-zero attractor set for a given parameter set.

#### Claim 1.

*Given the network structure in Fig. 1 of the main manuscript, the underlying dynamics in Eq. 2 of the main text, assuming 2 (semi-concave), with high killer T cell activity, can not contain two stable steady-states with non-zero high(low)-CAF and non-zero high(low)-immune cell components if the positive feedbacks involving Killer T cells-IL-2, and CAF-OPN can not yield more than one non-desert attractor*.

*Proof*. We prove this by contradiction. Let us suppose there exists a pair of non-zero, stable, steady states 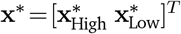 that is high (low) in both immune cells and CAF. From the Jacobian analysis of the dynamics, the total influence of CAFs on killer T cells (ℐ_*CAF*→*T*_) can be obtained as

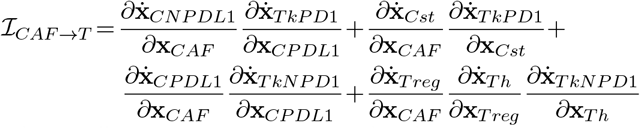

Further, the total influence of killer T cells on CAF (I_*T*→*CAF*_) can be obtained as

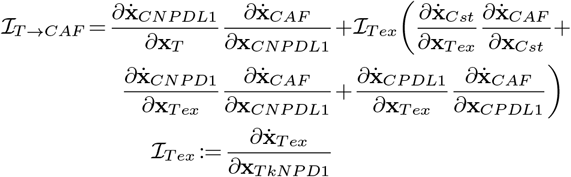

where **x**_*p*_ is the population of the node *p* in the node set.

Given the high activity of the killer T-cells and the Lipschitz assumption of 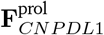 there exists an open interval 𝔹_Low*/*High_ ∈ℝ^*N*^ in the neighborhood of the steady states **x**^∗^, such that 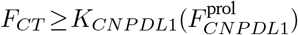, where *K*_*CNPDL*1_ is the Lipschitz constant for 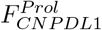. Similar is the scenario of tumor stem cells. Therefore, for high killer T-cell activity, there exists a point *p* ∈𝔹_Low*/*High_ with zero population for tumor stem cells and tumor cells without PDL1. This implies at *p*, although I_*CAF*→*T*_ remains negative, the net sign of ℐ_*T*→*CAF*_ becomes positive due to the vanishing of the terms 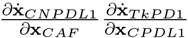 and 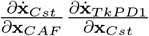.

For the existence of two stable, steady states, there needs to be at least one point **x**_*ext*_ on the plane 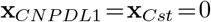 with 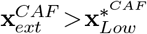 and 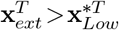 such that both the killer T cell and CAF components of the vector field evaluated at **x**_*ext*_ is positive even when **x**_*ext*_ is exterior to the domain of attraction of 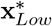 which is not the case owing to the mutually opposite net signs of ℐ_*T*→*CAF*_ and ℐ_*CAF*→*T*_. This contradicts the previous assertion.

#### Remark 3.

*The existence of two mutually opposite steady states becomes viable if the underlying dynamics constitute a competitive system between the CAF and the killer T cells [2]. Further, from the definition of competitive systems, the following conditions need to be satisfied*

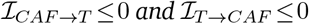

*where*, ℐ_*p*→*q*_ *represents the net influence of node p ∈ X on node q ∈ X. As seen in Claim 1, in the presence of exhausted T cells and hyperactive killer T cells, CAF and the killer T cells effectively engage in a negative feedback loop (Fig.**??**(a)) with a reinforcing net influence from the killer T to CAF— This violates the condition for competitive systems. Therefore, TMEs without the exhausted T-cells and moderately active killer T cells may exhibit both immune-rich and fibro-rich behavior*.

Therefore, we show that under Assumptions 1-3, the cardinality of the non-desert attractor set is less than equal to one for a given parameter set.

### 5 Response to ICI treatment

This section investigates the response of different TME subtypes to anti-PD1 treatment. We begin with the immune desert TME phenotype.

#### 5.1 No killer T cells, no response to ICI

##### Proposition 2.

*The immune desert TME subtype for the proposed TME in Fig. 1 of the main manuscript and for the dynamics in (1) does not change the tumor cell population when subjected to anti-PD1 therapy*.

*Proof*. As discussed in the methodology section, the system should be *controllable* by the external input *F* for the TME to be responsive to ICI treatment. As discussed in Proposition 1, the steady-state population of killer T cells in the immune desert condition is zero. Given the dynamics in Remark 4, and Eq.2, the linearized system matrices 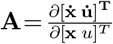 and 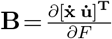 evaluated around the immune desert TME adopts the following structure

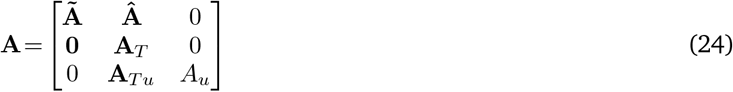

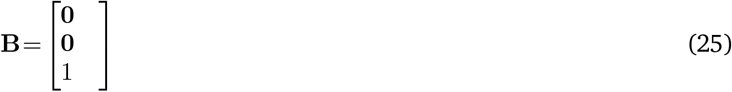

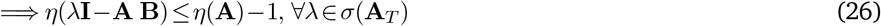

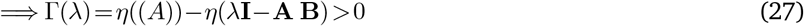

where, *η* is the rank operator. Therefore, the system is not controllable by *F*, indicating non-responsiveness to anti-PD1 therapy.

#### 5.2 Fibro-desert: Favourable scenario for ICI

We first start with the pre-ICI scenario of fibro-desert (Non-immune-desert) scenario.

##### Remark 4.

*We show that under certain restrictive conditions on the killer T cell activity and assumptions underlying the exhaustion rate of PD1^+^ T cell in the presence of PDL1^+^ tumor cells, the fibro desert TME does exhibit a unique stable, steady state— This does not eliminate the options for oscillation. We here lay out the concrete conditions for a unique stable steady state under the fibro desert regime in pre-ICI conditions.*

*Let us denote the population of three tumor cell states x_1_ (stem), x_2_ (PDL1^−^), and x_3_ (PDL1^+^), and killer T cells x_4_, exhausted T cells x_5_ and. 𝒮ince the proliferation of TAM is heavily dependent on the CAF population, we ignore the effect of TAM in the fibro desert region [3]. We also argue that including the effects of TAM will not alter the following result. Therefore, the approximated dynamical system around a fibro desert TME can be expressed as*

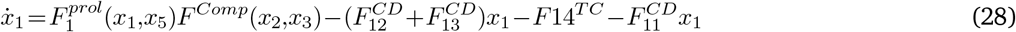

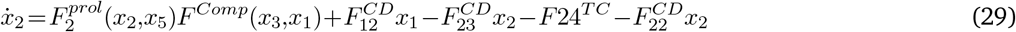

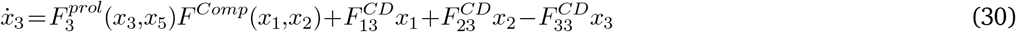

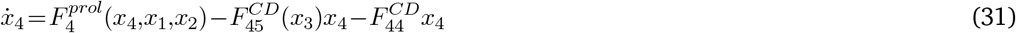

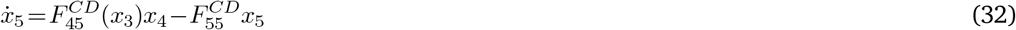

*Under high T cell activity and non-immune desert conditions, 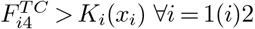, where, K_i_ is the Lipschitz constant for the proliferation rates 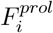− This implies the steady states of the stem and the PDL1^−^ tumor cells shall be zero. Further, let us assume a set of isolated steady states (as denoted by **x**^∗^) in the positive orthant exists for the system, excluding the trivial steady state of origin. It is to be noted that due to the non-immune desert condition, and due to 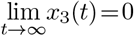being an unstable solution, there exists no line segment from two intermediate points in **x**^∗^ that passes through the plane x_3_ =0 or x_4_ =0.*

*Additionally, we assume the proliferation rate* 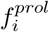 *can be factorized as* 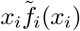. *Further, we also assume* 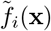 *in Eq. 4 is a monotonically decreasing function of x_i_. At each point in **x**^∗^, the expression* 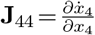 *can be evaluated as*

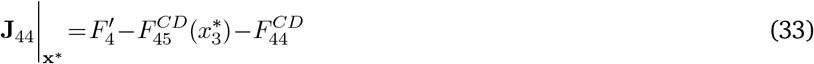

*where* 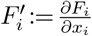 *Due to zero steady state values for x_1_ and x_2_, J_44_ can be written as*

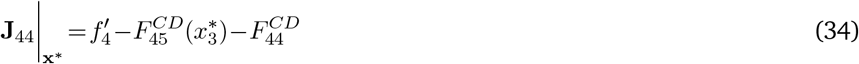

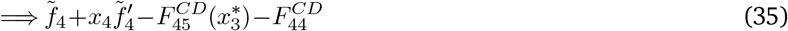

*Now, since **x**^∗^ only contains the steady states*

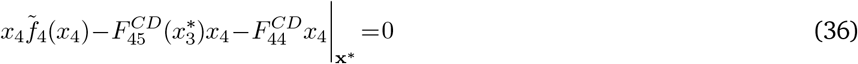

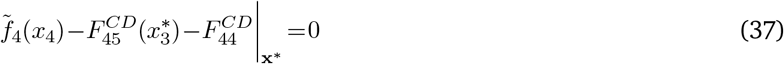

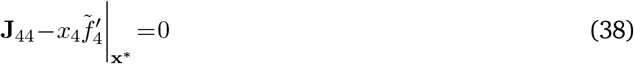

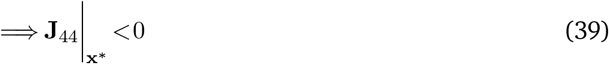

*similarly*, 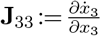 *can be written as*

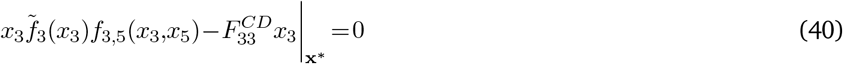

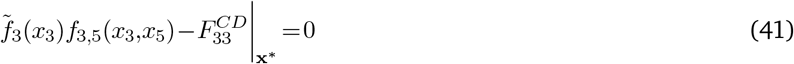

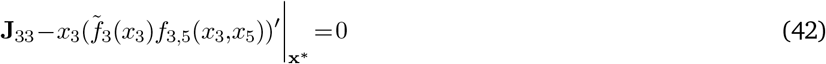

*As it can be seen from (42)*, 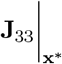 *is negative if and only if the product function f_3,5_(x_3_,x_5_) adopts a non-positive slope with respect to x_3_ at all **x**^∗^— this is denoted as the negativity condition*.

*Additionally, it is easily verifiable from (32) 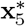 is a function of 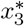 and 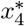. Therefore, if the negativity condition is satisfied, then all the Jacobian of the system centered around all the points in **x**^∗^ contain negative diagonal elements. Further, all the loops in the system consist of a negative sign— this implies all the points in **x**^∗^ are stable. since there can never exist two stable, steady state points adjacent to each other in a well-posed dynamical system (the solutions **x**_3_ =0 or **x**_4_ =0 do not lie in between), the only possibility is **x**^∗^ is singleton— The fibro desert TME only under aforementioned conditions adopt a unique stable, steady state.*

The fibro-desert, in a pre-ICI setting, TME may lead to oscillation, or in the specific scenario described in Remark 4, the pre-ICI population of tumor cells is sensitive to the oncogenic role of the exhausted T cells. However, an effective anti-PD1 therapy eliminates the dependence on the oncogenic role of the exhausted T cells.

##### Proposition 3.

*Under assumptions 1, 2(semi-concave), 3(high-killer T cell activity), and high killer T cell proliferation, the dynamical system in (1) underlying a fibro desert TME converges to a unique (zero), stable, steady state when subjected to effective anti-PD1 therapy*.

*Proof*. As it can be seen from the network in Fig. 1, under non-immune desert conditions, the immune module cannot contribute to the multi-stability of the underlying dynamical system owing to the absence of loops in the immune module. Further, by effective anti-PD1 treatment, we mean that the removal of PD1 expression from the killer T cells happens at a rate faster than all the interactions in the network. Therefore, the anti-PD1 modified dynamical system (D) approximated around the fibro desert TME can be expressed as

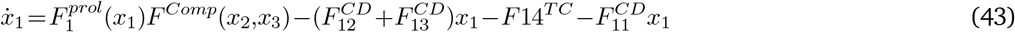

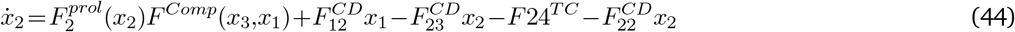

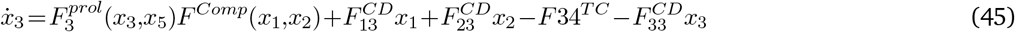

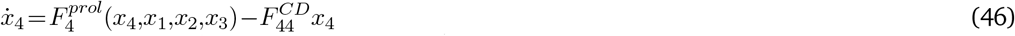

where, *x*_1_, *x*_2_, *x*_3_, *x*_4_ represents the population for stem, PDL1^−^, PDL1^+^ tumor cells, and PD1^−^ killer T cells respectively. The function 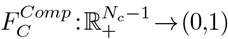 refers to the competitive inhibition exerted by different tumor cell states on each other.

Let us define another dynamical system 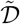

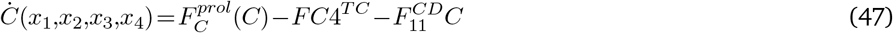

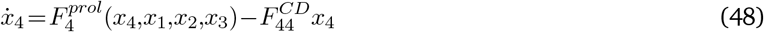

where, 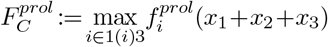.Due to the assumption of high killer T cell activity and non-immune desert environment,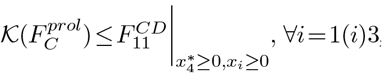, equality holds when *x*_4_ or *x*_*i*_ is zero. 𝒦 is the Lipschitz constant of 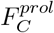– This implies the steady state solution for *C*(*t*) is zero. Since,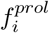 is semi-concave, 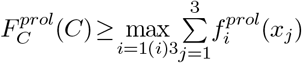 — This implies there exists an open neighbourhood 𝕆⊂ ℝ^3^ of *C*^∗^ = 0 such that ∀*C*(*t*) ≥ *x*_1_(*t*)+*x*_2_(*t*)+*x*_3_(*t*) ∈𝕆. Now, 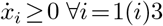 at *x*_*i*_ = 0 planes imply both 𝒟 and 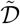 evolves in the positive orthant of the state space. Therefore, the only feasible solution in 𝒟 is for all the tumor cells to go to zero at large times. This concludes the proof.

#### 5.3 Immune-rich: From reduction to complete removal

The pre-ICI attractor set corresponds to an immune-rich TME subtype, providing a favorable response to the ICI response *i.e*, the post-ICI attractor has either no or lesser tumor cell count than its pre-ICI counterpart. According to our analysis, whether ICI therapy leads to completely eliminating or reducing tumor cells depends on numerous factors, such as the high proliferation rate of the tumor cells and low T cell activity. Here, we propose a sufficient condition for the reduction scenario that sets it apart from the trivial scenario.

##### Remark 5.

*Given the dynamics of the TME network in Fig. 1 of the main manuscript and the underlying dynamics in (1) satisfying assumptions 1-4, the ICI therapy cannot lead to complete tumor cell elimination if the following condition is satisfied*

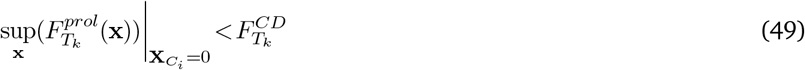

*where* 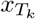, *and* 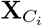 *denote the killer T cells and different tumor cell states, respectively. K is the lowest Lipschitz constant for the growth and proliferation term 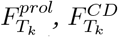represents the conversion from and death term of killer T cells*.

*(49) leads to the scaneario 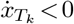. Therefore, throughout the plane 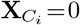, the killer T cell count continues to reduce, rendering the existence of an attractor with a non-zero killer T cell component and zero tumor cells infeasible. Further, due to the high growth rate of the tumor cells, all the steady states with zero killer T cells and zero tumor cell components also remain unstable. Therefore, complete elimination is not possible in this scenario*.

*Further, if ∃ an ϵ>0 such that 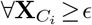, the following is satisfied*,

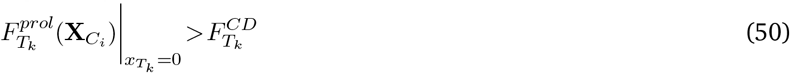

*As evident from (50), if the tumor cell component corresponding to the steady states with zero killer T cells (and non zero tumor cells) is greater than or equal to the ϵ barrier, the immune-desert scenario can be avoided*.

*Overall, if both the conditions hold true, then complete elimination becomes infeasible. At the same time, (50) along with the condition thereof ensures the infeasibility of an immune-desert-like attractor— This translates to the possibility of a post-ICI scenario with reduced tumor cell population (which is still ≥ϵ) and significant killer T cell population*.

#### 5.4 Immune accessibility index

We did not assume an expression of the immune accessibility index throughout the theoretical derivations in this work. We formally define the immune accessibility index *I* as the ratio between the immune-accessible (IA) tumor cells and fibroblast-protected tumor cells.

Further, for the purpose of simulation, we propose the following way to calculate the immune accessibility index (*I*)

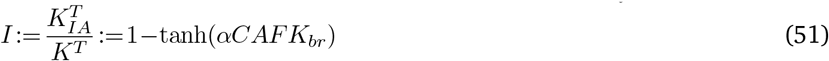

where, *α* determines the fraction of CAFs in close proximity to the tumor cells, *K*_*br*_ is the barrier formation rate for the CAFs, and 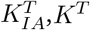 are the carrying capacities of the tumor cell states accessible by the immune system and the same before the compartmentalization by the CAFs.

As depicted in (51), *I* is a bounded quantity in the closed interval [0,1] wherein *I* =1 refers to the situation where all the tumor cells are accessible to the immune system, whereas *I* =0 denotes the impenetrability of the immune cells to the TME.

Therefore, with the introduction of the immune accessibility index, we characterize the immune-rich TMEs as the scenario where 1. the immune dessert is not in a stable, steady-state and 2. the immune accessibility index is high. On the other hand, the fibro-desert scenario is identified with 1. fibro-desert being an unstable steady state and 2. low immune accessibility index.

As established before, the CAF population around the tumor cell states significantly affects the immune accessibility. Therefore, for any given *CAF* population, *α* and *K*_*br*_, we propose the following model for the penetration of the immune cells as

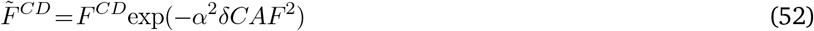

where *δ* is the width of CAF barrier.

It is evident from (52) that for large *CAF* population and non-zero *α*, the term *F* ^*CD*^ goes to zero rapidly. Therefore, the following remarks characterize the fibro-rich behavior.

##### Remark 6.

*For the network dynamics in* (1), *according to the condition posed for the responsiveness to ICI treatment, (a) the system in (1) has to be controllable to anti-PD1 species (u(t)) and (b) the quantities 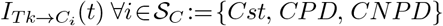 are non-positive everywhere ∀ t≥0 and non-zero at least once*.

*In the presence of a CAF barrier, let us denote the population immune accessible (IA) and CAF-protected (FP) tumor cell states* 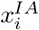 and 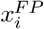 *∀I* ∈𝒮_*C*_ *respectively. Therefore, the common forward paths from the killer T cells to tumor cells across different states in a compartment is*

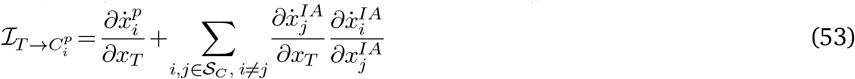

*where, p ∈ {IA, FP }. Now, for CAF-protected tumor cells, the term* 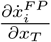 *becomes extremely small. The sign of the element* 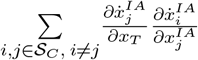 *is non-negative due to the competitive relation between the immune-accessible and CAF-protected tumor cell states. Further, in the fibro-rich scenario and low immune index, the first term* 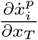 *for is almost zero. On the other hand, for the CAF-protected tumor cells, the sign of the second term is always non-negative. Therefore, the overall sign of* 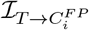 *is non-negative almost everywhere. This implies that for fibro-rich scenarios and low immune accessible index (I), the CAF-protected tumor cell population increases post-ICI*.

### 6 IL2-based treatment for immune-desert TME

As established in Proposition 2, the immune desert phenotype does not yield any significant response to ICI-based therapies. This necessitates additional perturbation strategies in conjunction with traditional ICI therapy. It is to be noted that the non-response to ICI therapy happens due to the absence of reachability from the anti-PD1 drug to the tumor cells. Further, the absence of reachability is attributed to the depletion of killer T cells. Therefore, the only way to circumvent the immune desert condition is to ensure the existence of at least one non-immune-desert attractor.

Additionally, a mere guarantee of a non-immune-desert attractor does not lead to a no-response for any initial conditions sufficiently close to an immune-desert attractor, which may still converge to zero killer T cell population. Therefore, there exist two possible way outs: 1) destabilization of immune-desert attractor or 2) driving the state trajectory away from the domain of attraction of immune-desert attractors. Further, it is to be observed from the TME network in Fig. 1 of the main manuscript that the only controllable species governing the proliferation of killer T cells in IL-2. Further, the dynamics of Killer T cells and IL-2 can be represented as

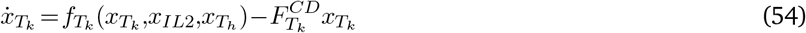

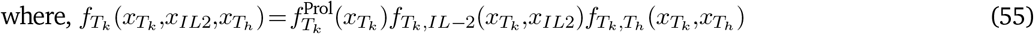

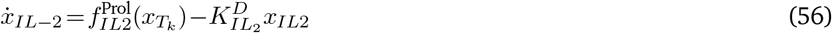

It can be easily verified that the for 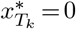 leads to zero final IL-2 level. For the immune desert condition, according to Proposition 1, 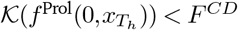 where, 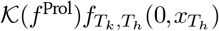 is the Lipschitz constant with respect to 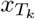. Let us define a set 𝒮 :{*x*_*IL*2_ ∈ℝ such that 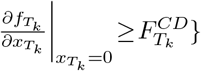. We are now ready to present the following result.

#### Proposition 4.

*For the given network in Fig. 1, if the set* 𝒮^*IL*−2^ *is non-empty there exists a threshold constant supply rate* 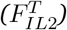 *of IL-2 that starts before the anti-PD1 treatment such that the immune dessert TME subtype becomes unstable*

*Proof*. Observing the condition in the given statement does not eliminate the possibility of an immune desert condition. The proof is constructive. With an external supply rate of *F*_*IL*2_, the dynamics of the concentration of IL-2 (*x*_*IL*2_) can be expressed as

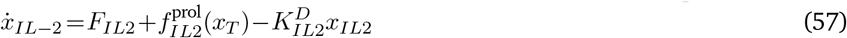

In immune-dessert condition, the steady state of *IL*2 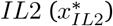 becomes

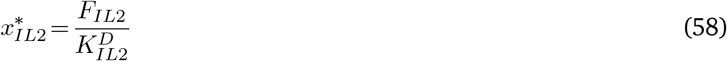

Now, it has already been shown in the proof of Proposition 1 that the systems matrix linearized around the immune desert steady state can be expressed as upper triangular matrix. In fact, the sub-matrix 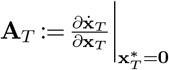 is also diagonal (or upper triangular for immune cold subtype). Therefore, in the pre-ICI condition

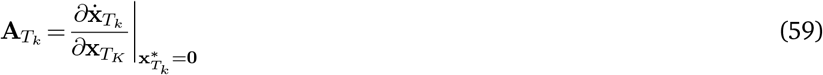

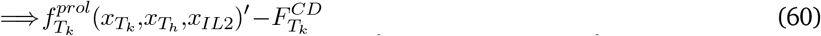

Due to non-zero 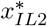 owing to *F*_*IL*2_ (60) can be re-written assuming the dynamical form given in Eq. 4 of the main manuscript,

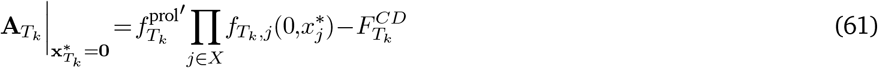

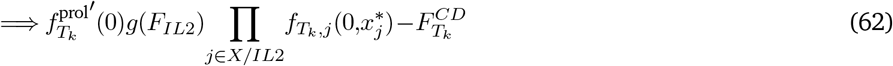

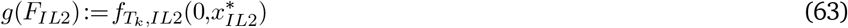

Further, since the set 𝒮^*IL*−2^ is non-empty, there exists an IL-2 level 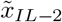 such that the expression in (62) adopts a strictly positive value. Further, as evident from (62), the immune-desert condition becomes unstable if 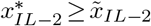.From the Archimedean property of ℝ^+^, the set 𝕊 := *n* such that 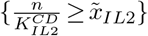 is non-empty. Choosing *F*_*Il*2_ :=inf 𝕊 completes the proof. Note the least upper bound property of ℝ guarantees the existence of *F*_*IL*2_.

Proposition 4 requires a constant supply rate of IL-2, which may not be practically feasible owing to the well-known toxicities of IL-2. Therefore, we explore the second option. Note that since function 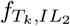 is a strictly monotone increasing of *IL*−2, for a given 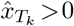, if there exists an IL-2 level 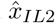 such that 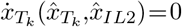, then 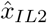 is unique. Further, let us also define a set 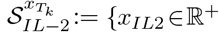 such that 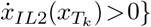. Note the set 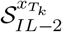 is always bounded above (from Theorem 1). Therefore, sup 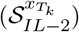 exists.

This inspires the following proposition

#### Proposition 5.

*If the desert condition in Proposition 1 is satisfied and if there exists an open interval 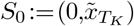, such that 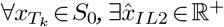,such that 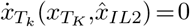 and if sup 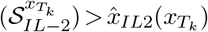 for at least one 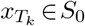 then the dynamical system in (1) either contains another non-desert stable steady state or stable limit circle*.

*Proof*. Since, there exists an open interval *S*_0_ in 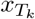 such that 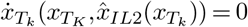, any point 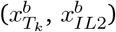 above the curve 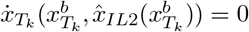 should have 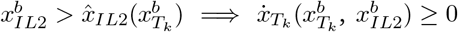. Similarly, for any point 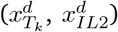 below 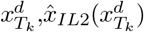 has 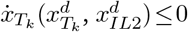. Since the desert condition as laid out in Proposition 1 is satisfied, the origin is a locally stable steady state. Therefore, there exists an open interval ℬ⊂*S*_0_ in the neighborhood of origin such that 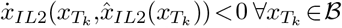.Therefore, for the origin to be globally stable steady state 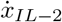 has to be non-positive along the curve 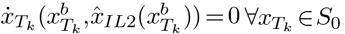. A violation of this condition implies a sign change of 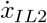 along the curve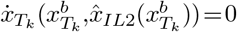 for non-zero positive *x*_*IL*2_ indicating the possibility of another steady state for the well-posed dynamical system.

Since, there exists at least one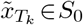 such that 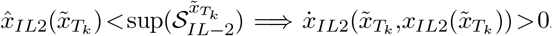. This violates the unique global stability of the origin. Further, Theorem 1 guarantees a positive invariance set in ℝ^+^ ∪0. This guarantees that all the trajectories starting from 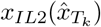 either converge to a non-desert steady state or enter into a stable limit circle with a strictly positive killer T cell component. This concludes the proof.

#### Lemma 7.

*If the conditions for Proposition 5 are satisfied, then there exists a threshold one-time IL-2 injection level such that the solution to (1) converges to a non-immune dessert steady state or limit circle if the initial killer T cell population 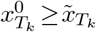 and* 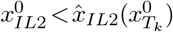

*Proof*. If conditions for Proposition 5 are satisfied, then there exists a set ℬ^*C*^ ⊂ℝ ^2+^ such that for all the initial conditions started in ℬ^*C*^ does not converge to zero. Further, from Proposition 5, the solutions with the initial condition 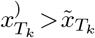 and 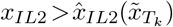 do not return to the origin. Therefore, for a given initial T-cell population 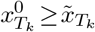, the amount of one time IL-2 injection required to drive the state trajectories out of origin is given by 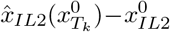.

#### Remark 7.

*Without the removal of the PD1 program, the multiplier term 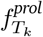decreases. Additionally, owing to the pre-ICI T-cell exhaustion, the second term 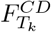 increases. Therefore, the amount of additional IL−2 required in pre-ICI to destabilize the immune desert is higher than its post-ICI counterpart*.

*Similarly, due to the reasons stated above the domain of attraction for the origin in pre-ICI setting is significantly larger than the same post-ICI therapy i. e., 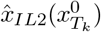 increases. This implies that it requires a larger (than the post-ICI equivalent) one-time IL-2 intake in pre-ICI conditions to steer the state trajectories towards an immune-hot steady state*.

### 7 OPN knockout: Transition to fibro-desert

The fibro-rich scenario, as defined in the main manuscript, can be identified with a low immune accessibility index and non-fibro-desert attractors/ limit circles. Due to the abundance of fibroblasts, immune accessibility is poor. Therefore, the post-ICI tumor cell population does not undergo any significant deviation from its pre-ICI counterpart. Since the immune accessibility index (*I*_*a*_) is a strictly monotonically decreasing function of the CAF population, improving *I*_*a*_ amounts to reduce the CAF population significantly. Further, the CAF population growth primarily depends on the autocrine reaction through OPN. Therefore, we select OPN as the potential therapy target to modulate the proliferation rate of CAF such that, in the best-case scenario, the TME transitions into a fibro-desert scenario.

Given the dynamics of CAF and OPN population (*x*_*CAF*_, *x*_*OPN*_, respectively)

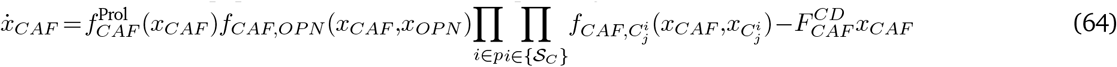

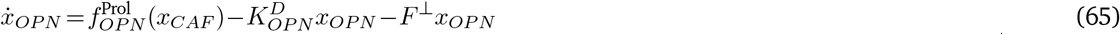

where, *p* :{IA, FP} denotes the set for immune accessible and CAF-protected tumor cells. Further, 𝒮_*C*_ :{*CST, PDL*1^+^, *PDL*1^−^} denotes the stem, PDL1^+^, and PDL1^−^ tumor cells. *F* ^⊥^ refers to the externally controlled OPN clearance rate.

We propose the following condition:

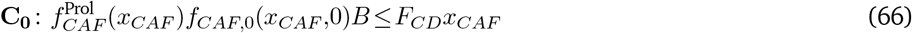

where *B* is the upper bound of 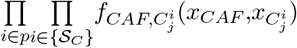. Note that the Assumption of boundedness guarantees the existence of finite *B*.

#### Abuse of notation

We define a pseudo-variable 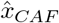 with the dynamics as following

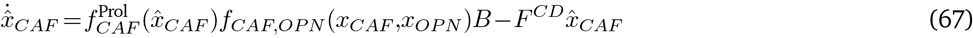

From Lemma 2, 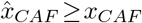. We shall work with 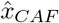 from this point onwards. Therefore, we take the liberty to drop the symbol ‘^’ over the pseudo-variable 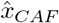

We propose the following proposition

##### Proposition 6.

*If the condition **C**_0_ is satisfied, and there exists a non-fibro-desert attractor for the dynamical system in (1), then there exists a threshold OPN clearance rate F^⊥^ such that the fibro-rich attractor ceases to exist and the fibro-desert steady state emerges as the only attractor*.

*Proof*. Since, **C**_0_ is satisfied, for *x*_*OPN*_ = 0 and 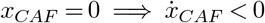. Further, for any *x*_*CAF*_ ∈ 𝒩 (0,*r*) ∃ a δ *>* 0 for which 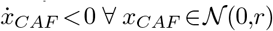 and *x*_*OPN*_ *<*δ(*x*_*CAF*_). This establishes the local stability of the fibro-desert-like attractor when the OPN level is zero. We now show that the non-fibro-desert attractor withers away for an OPN level below a critical value.

Theorem 1 guarantees a positive invariant set in the state space. Denote the maximum boundary value for *x*_*CAF*_ as 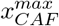.

Similarly, for *x*_*OPN*_, it is 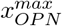 We define the set 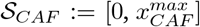, and 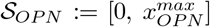. Further, consider the set 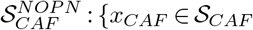, such that 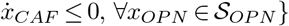. Note 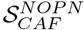 contains the point *x*_*CAF*_ = 0. Therefore, the set 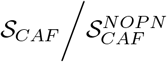 is defined as

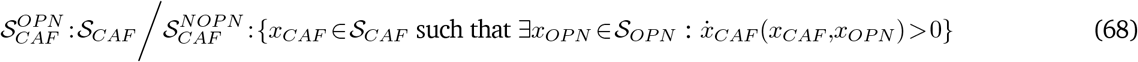

Note, 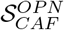 cannot be an empty set. If that is the scenario, then 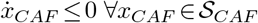 and *x*_*OPN*_ ∈ 𝒮_*OPN*_. Therefore, there is no sign change in 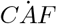 throughout the state space, ruling out multiple stable steady states for CAF. Therefore, the only attractor in the system has fibro-desert properties, which violates the proposition’s initial premise, which is the existence of a fibro-rich attractor with non-zero OPN concentration.

Denote the set 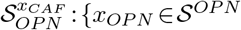 such that 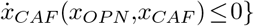. Since, for a given *x*_*CAF*_, *f*_*CAF,OPN*_ is a monotone increasing function of *x*_*OPN*_ and condition **C**_0_ is satisfied, then sup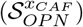 exists and is greater than zero. Therefore, all the elements of the set 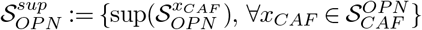 is non-zero and finite. Therefore the quantity 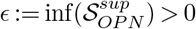. Therefore, if 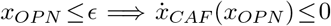 throughout the set s_*CAF*_ with the only stable steady state with zero CAF component. The equality holds only at *x*_*CAF*_ =0. Therefore, the stable non-fibro-desert attractor ceases to exist.

Further, we the following propose sufficient condition for the OPN clearance rate to regulate the OPN level below *ϵ*

This proves the existence of an OPN clearance rate such that the Fibro-desert steady state remains the only attractor throughout the state space. This concludes the proof.

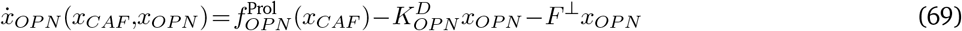

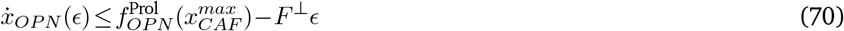

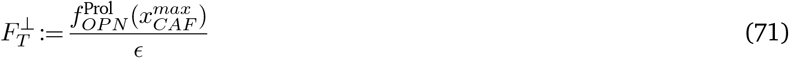

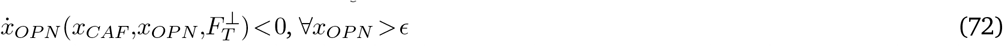

Proposition 6 assumes **C**_0_, which ensures the feasibility of a fibro-desert attractor. Even if the condition **C**_0_ is not satisfied, the CAF population post OPN knock-out is significantly reduced compared to the pre-knockout scenario. This is immediate from Lemma2. Further, it is implicit in **C**_0_ that the population of wild-type fibroblasts is non-significant in the analysis of fibro-rich TME subtypes. This is because due to the high conversion rate from the wild type to the invasive CAFs and low natural proliferation rate, the fibro-rich-like attractor remains depleted of wild-type fibroblasts. Therefore, even in the post-knockout scenario, the conversion term 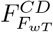 is assumed to be zero in this case. [3, 3–6, 6–20] [10, 12, 19, 21–74].

